# Diffusion of kinesin motors on cargo can enhance binding and run lengths during intracellular transport

**DOI:** 10.1101/686147

**Authors:** Matthew Bovyn, Babu Reddy, Steven Gross, Jun Allard

**Author notes:** Center for Complex Biological Systems, University of California, Irvine (Matthew Bovyn, Steven Gross, Jun Allard).

## Abstract

Cellular cargos, including lipid droplets and mitochondria, are transported along microtubules using molecular motors such as kinesins. Many experimental and computational studies of cargos with rigidly attached motors, in contrast to many biological cargos that have lipid surfaces that may allow surface mobility of motors. We extend a mechanochemical 3D computational model by adding coupled-viscosity effects to compare different motor arrangements and mobilities. We show that organizational changes can optimize for different objectives: Cargos with clustered motors are transported efficiently, but are slow to bind to microtubules, whereas those with motors dispersed rigidly on their surface bind microtubules quickly, but are transported inefficiently. Finally, cargos with freely-diffusing motors have both fast binding and efficient transport, although less efficient than clustered motors. These results suggest that experimentally observed changes in motor organization may be a control point for transport.

## 1 Introduction

To establish and maintain their internal organization, eukaryotic cells employ molecular motors in the kinesin, dynein and myosin superfamilies to transport organelles and other cargo along microtubule (MT) and actin filaments. Cargo transport is a complex, multi-step process at both cell-scale [1, 2] and cargo-scale [3, 4]. One key step that has been less studied so far is initiation of transport, specifically, how long it takes to attach a cargo to the MT in the first place. *In vitro*, we recently reported that the time for a cargo to reattach to the MT after detaching in an optical trap is on the order of 1 second at viscosities 1x-10x water [5], and depends strongly on the size of the cargo, indicating that cargo rotation plays an significant role. How does this extrapolate to the cytoplasmic environment, where viscosities are estimated to be orders of magnitude larger (see 4.2)? We begin with a simple calculation, presented in Fig 1A. We ask how long it would take for a motor initially located opposite the MT to come within range of the MT, assuming a motor (or cluster of motors) is rigidly bound to a cargo. For cytoplasic viscosity estimates (gray line), for a cargo of 500nm-1000nm in diameter, we estimate the time to be 10^2^ – 10^3^s. (For details on the calculation see section 4.5. Note that, since the times for motor repositioning are so slow, the choice of local motor attachment rate value *k_on_* ≥ 5s^−1^ [5] does not significantly affect total cargo attachment time.) Yet, in *vivo*, in COS1 cells, after detachment in an optical trap [6], rebinding times were ~10 s for dynein-driven transport and ~7s for kinesin-driven transport ([6] and section 4.4). In an even more extreme discrepancy, binding times for the lipid droplets purified and measured in *vitro* were found to be 0.2 s ([6] and section 4.4).

**Figure 1:**
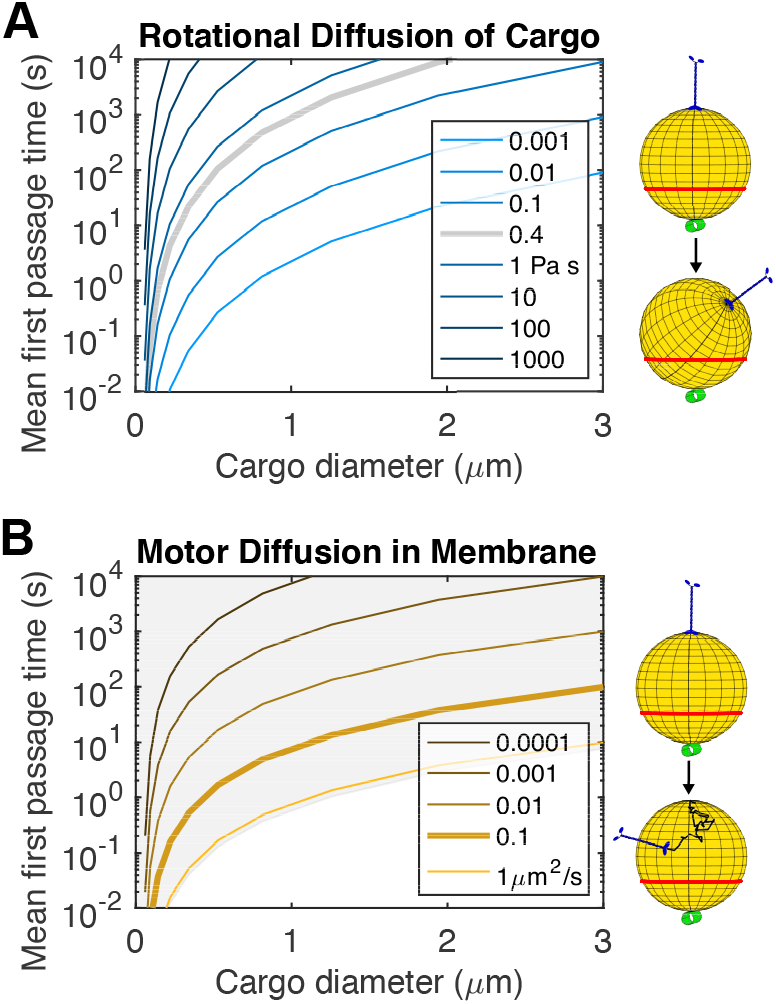
A motor (blue in right schematic) initially located opposite a microtubule (green) must move (to below red line) in order to be bind the microtubule. Mean first passage times (MFPTs) for this movement is shown for either: **A**: rotational diffusion of the cargo. Shown for various values of viscosity of the surrounding fluid. Values are chosen to span the range of values estimated for cytoplasm. Grey curve is at our best guess for viscosity experienced by intracellular cargos, see 4.2. **B**: diffusion of the motor in the cargo surface. Shown at various values of the diffusion coefficient. MFPTs for diffusion coefficients achievable when the surrounding fluid has viscosity of 0.4 Pas are shaded in grey, see 4.6. For details on calculation, see 4.5. Horizontal lines show the range of experimentally measured values.

What accounts for this discrepancy? One possible explanation is a difference of concepts of attachment rate. There are at least four [5]: The time for a cargo, on first approach to a MT (e.g., from the organelle that generated it), the time to rebind after pulling out of a trap, the concentration-dependent local attachment rates of the motor head to the MT lattice, and the local attachment rate of a second motor to the MT on a multi-motor cargo. Another possibility, related to the first and the focus of this work, concerns mobility of the motors on the cargo. Many cellular cargos are bounded by lipids, raising the possibility that motors diffuse in those lipids. Freedom of motors on the surface of cargos has been recently studied. In the actin-myosin transport system, both the velocity and run-lengths of lipid vesicles has been shown to depend on membrane composition [7]. Several papers have compared the collective action of kinesin motors which are free to diffuse in membranes to cases where they are rigidly attached to a surface, both in MT gliding assays [8] and bead assays [9]. These studies show that motor freedom in membranes can influence transport properties, but it is unclear how to translate the *in vitro* results into the context of the cell, especially since transport is slowed in the gliding assay, while it is sped up in the bead assay. Furthermore, Li et al. [9] find that, when motors are allowed to diffuse, there is a small increase in velocity, with no difference in how far cargos travel overall. Thus, the impact which freedom to diffuse may have on cargo transport in the cell is unclear.

As a first step, in Fig 1B we perform the equivalent of the rigid rotation calculation in Fig 1A, now assuming motors are free to diffuse in a fluid membrane. For diffusion coefficients typical of membrane-anchored proteins (thick line, BioNumbers BNID:114189 [10]), for a cargo of 500nm-1000nm in diameter, we estimate the mean time for a motor to diffuse, from far from the MT to within binding radius, to be 1-10s. Therefore, the diffusion is fast enough that it might help accelerate motor binding; membrane fluidity might in principle be a key factor in cargo binding and transport initiation. However, the calculation shown in Fig 1B is highly simplified. It ignores binding times, motor on rates, and does not address subsequent transport.

Another possible resolution is that many motors are dispersed around the surface, reducing the mean angle through which the cargo must rotate before one of the motors is within reach of the MT. Cellular cargos often have several motors associated with them [11, 12]. However, several motors often engage the MT simultaneously [6, 13], and rigidly dispersed motors are unlikely to be able to bind simultaneously, and less able to form clusters once the cargo is bound to the MT [4].

In this paper, we carry out a comprehensive computational study of how cargos with different motor organizations are transported with a focus on the less-explored case of cargos which have motors that are free to diffuse in the membrane. This is particularly relevant since some cargos appear to switch motor distribution as they progress through their life cycle [3]. This requires us to compare transport properties over a wide range of membrane fluidities, from fluid to solid. We construct a computational model of cargo transport which includes multiple motors that are able to diffuse in the cargo membrane. While there is a rich history of theoretical studies of motor-based transport [14, 15], the ability to describe the full range of membrane fluidity represents a significant advance, including over 3D models with rigidly attached motors [4, 16]. Since there are two viscosities in the system (the membrane and the cytoplasm), forces translate into motion in a non-trivial, coupled manner. Furthermore, the two viscosities induce diffusion (thermal fluctuations) with different amplitudes, so motors experience statistically correlated noise. Overcoming these computational challenges required that we separately formalize the stochastic equations of motion for all motors and cargo, analytically manipulate these in full generality into a frame of reference in which the motion can be simulated using an Euler-Maruyama scheme, and combining this with the chemomechanics of motor binding and unbinding. This allowed the computationally efficient simulation needed for the large parameter sweeps we report, in contrast to lookup-table approaches or numerical iteration methods. (Several recent models have included motors that rearrange on the cargo [17, 18]; however, they limited their studies to parameter regimes in which the complicating factors above were absent.) This software is openly available at github.com/mbovyn/Motor-Cargo-Simulator (DOI:10.5281/zenodo.4325111). The results of our computational model reveal that the changes in motor organization and mobility have significant impact on MT binding, transport and force generation. We made a particular effort to simulate behavior in viscosities relevant to the cellular environment. Our work points to parts of cargo transport that are not well understood, and informs the picture of overall transport.

## 2 Results

### 2.1 A computational model of cargo transport including freedom of motors to diffuse in the cargo membrane

We wish to explore the impact of different classes of surface organization of motors on the transport steps of binding and running (Fig 2A). We choose four extreme cases, as shown in Fig 2B, which we term organization modes. The first two modes have motors bound rigidly to the cargo. They differ in how motors are spaced; the first mode we term “rigid clustered” places all the motor anchors at the same point, while in the second mode, which we term “rigid dispersed”, the motor locations are random, drawn from a uniform distribution over the surface. The other two modes have motors which are free to diffuse in the cargo surface. In the “free independent” mode, the motor anchors do not interact with each other, other than through forces they exert on the cargo. In the final mode, termed “free clustered”, all motor anchors are bound together, but the ensemble is able to diffuse in the cargo membrane. In this mode, we assume the cluster of motors diffuses with a reduced diffusion coefficient so that 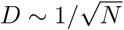 where *N* is the number of motors, which is consistent with motors being arranged in roughly a disk (rather than, e.g., in a row, see 4.7).

**Figure 2:**
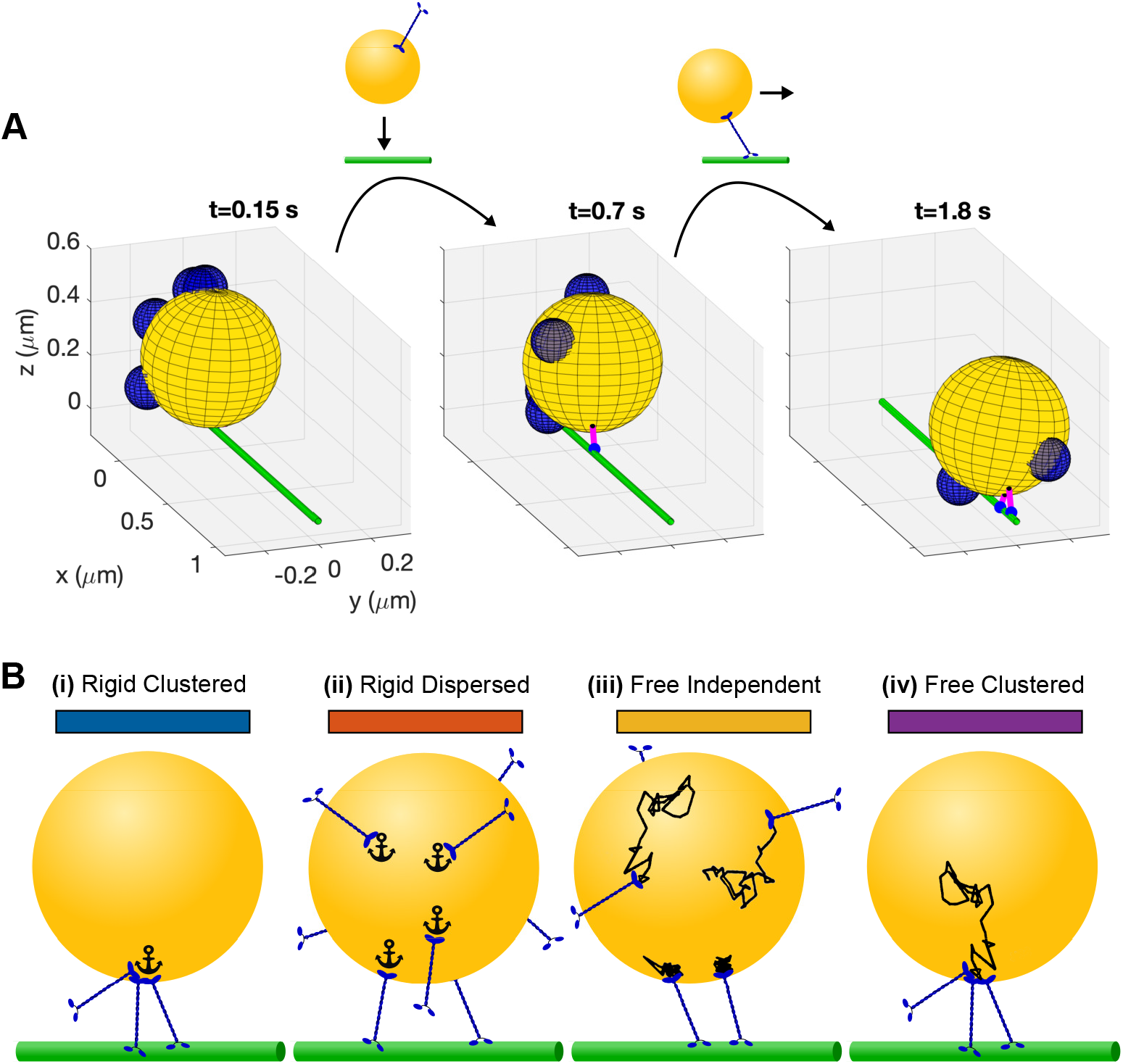
Transport outcomes and motor organization modes to be investigated. **A**: We separate the cargo transport process into binding and run steps. Before binding, first bound, and transported states are represented by simulation snapshots. The cargo is yellow and the microtubule is green. The blue hemispheres represent the reach length of unbound motors, which have their anchor point at the center of the hemisphere. Bound motors are represented with a magenta stalk, small black anchor point and larger blue sphere at the center of mass location of the motor heads. **B**: Motor anchoring modes to be investigated. Cargo shown in yellow, motors in blue and microtubule in green. We name the states: rigid clustered (**i**), rigid dispersed (**ii**), free independent (**iii**), and free clustered (**iv**). Solid color bars indicate the color used for this mode in the rest of the text.

To investigate transport outcomes of cargos in these four different organization modes, we construct a three dimensional model of a cargo, the motors, and a MT. This model includes properties of the cargo, such as size and shape, properties of the motors, such as length and binding rate, and properties of the environment, such as viscosity. For every property we estimate ranges and typical values for cargos in the cell. Descriptions of how these values were estimated and relevant information can be found in section A2.1. Values used can be found in table A1.

We model the cargo as an undeformable sphere. We attach motors to the cargo at points which we term the anchors. We model these motors using the well studied chemomechanics of kinesin-1. These motors can bind the MT when the anchor is within reach of the MT (blue hemispheres in Fig 2A represent motor reach length). Once bound, they step along the MT and unbind from it with rates that have been well-studied *in vitro*. We draw our chemomechanical model for kinesin from recent experimental [19] and modeling [20] efforts. For more details, see supplement sections A1.1.1 and A1.3.

As motors step, they exert forces on the cargo which would both tend to pull the cargo along through the surrounding fluid, and drag the anchor through the cargo membrane. In our model, forces which would drag the anchor through the cargo membrane result both in displacement of the anchor in the membrane and rotation of the cargo, in proportion to the anchor diffusion coefficient. Forces acting to move the cargo body do so against viscous drag, as we model the surrounding fluid as Newtonian. While recent measurements suggest the cytoplasm is an actively-driven, complex fluid with significant elasticity [21, 22], methods for simulating diffusion and the effect of further active forces in this environment are still in development. Modeling the cytoplasm surrounding the cargo as Newtonian allows us to qualitatively capture that some cargos in cells may be relatively free to move, while others may be significantly impeded by their local environment. For more information, see A2.1.3.

Both cargo and motors diffuse in their respective (3D or 2D) fluids. The cargo diffuses both rotationally and translationally with statistics governed by the Fluctuation-Dissipation Theorem, as implied by the viscosity of the surrounding fluid. Motors diffuse in the cargo surface with statistics governed by a diffusion coefficient. In general, complex movement may result from the interaction of motor anchors with different lipid domains [3] or other structures (for example, diffusion of cell surface proteins is influenced by the underlying actin cortex [23]).

A full description of the model and derivation of the equations we simulate can be found in supplemental section A1. The simulation framework is detailed in supplemental section A2 and parameter values listed in table A1 are used for all simulations, unless otherwise indicated.

To obtain transport properties of cargos in different organization modes, we simulate 100 or more stochastic trajectories and examine the resulting distributions. We simulate trajectories using a hybrid Euler-Maruyama-Gillespie scheme, and report a series of tests which show that the code reproduces expected results in some simplified situations in supplemental section A3. Snapshots from a single trajectory are shown in figure 2A. As time progresses, motors diffuse around the surface of the cargo, motors bind and unbind from the MT, and cargo orientation changes as it is pulled along by forces generated by the motors, as shown in video 1.

### 2.2 Cargos with free motors bind to the microtubule faster than those with rigidly anchored motors

For a cargo to be transported, one of its motors must first bind the MT. In this section, we use our model to investigate the time it takes for a cargo located near a MT to bind to it. Visualizations of example simulations for each organization mode are shown in videos 2–5.

We first compare the four organization modes as a function of the number of motors on the cargo, as Fig 3A shows for a 0.5 μm diameter cargo. Simulated cargos are allowed to diffuse rotationally, but not translationally, with no gap between the cargo and microtubule. Initial motor locations are picked from a uniform distribution over the surface of the cargo. We find that for cargos in the rigid clustered mode, the mean time to bind is long when there is a single motor, and stays constant as the number of motors on the cargo increases. This can be understood by considering the timescales in the problem; cargos in this mode spend most of their time waiting until the motors are near enough to the MT for them to bind, as rotation is slow compared to the characteristic binding time of a single motor. A cargo in the rigid dispersed mode with only one motor is identical to a rigid clustered cargo with one motor. For rigid dispersed cargos, however, we find the time to bind decreases drastically as more motors are added. The average angle though which these cargos must rotate before a motor comes within reach of the MT decreases as motors are added. This change is most drastic for the first few motors, with the time to bind of these 0.5 μm diameter cargos decreasing by an order of magnitude with the addition of only 5 motors.

**Figure 3:**
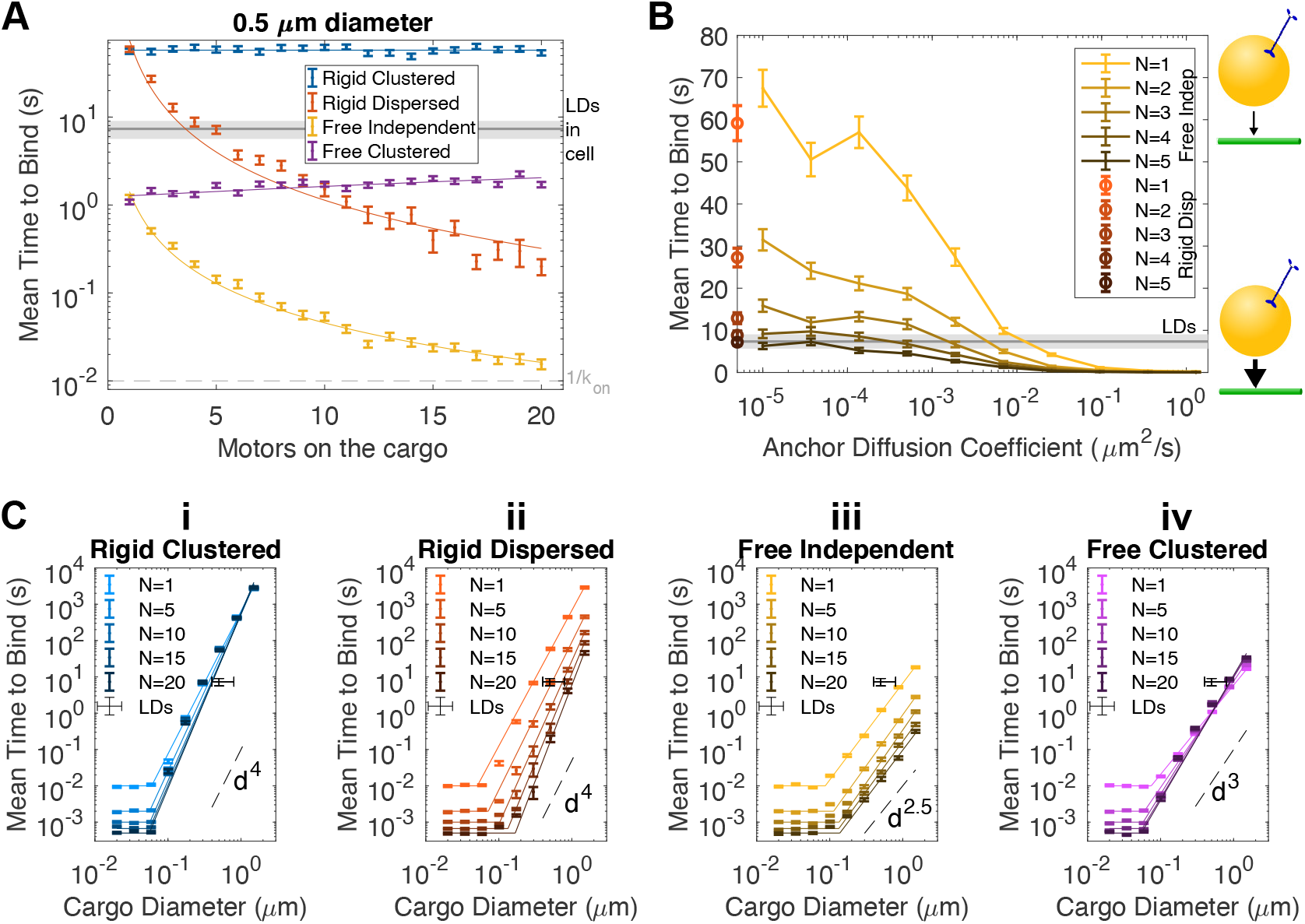
Times for cargos in different organization modes to bind to the microtubule Simulated cargos are allowed to diffuse rotationally, but not translationally, with no gap between the cargo and microtubule. Initial motor locations are picked from a uniform distribution over the surface of the cargo. In all panels, error bars represent standard error of the mean of 300 simulations. Underlying distributions are approximately exponential (see figure S2). The experimentally measured time for lipid droplets (LDs) to bind in the cell [6] is represented in each panel: Grey rectangles in panels **A** and **B** represent the measured mean +/- standard error of the mean. In **C**, additional error bars in diameter represent the approximate range of sizes of lipid droplets measured. **A**: Time for the cargo to bind to the microtubule for the four anchoring modes, as a function of the number of motors on the cargo. The characteristic time for a single motor to bind, assuming it is near the microtubule, is shown in dashed grey. Distributions of times to bind in the different modes are shown in figure S2A. Overlaid curves are fits, detailed in table S1. **B**: Dependance of the cargo binding time on the diffusion coefficient of the motor anchors in the cargo membrane for the free independent mode. Red error bars located on the left vertical axis are for the rigid dispersed mode. Distributions of binding times at the indicated diffusion coefficients are shown in figure S2Bi. Distributions of binding times for the lowest diffusion coefficient and rigid dispersed cargos are shown together in figure S2Bii. Lines between points are guides for the eye. **C**: Time for the cargo to bind as a function of the cargo radius for the four anchoring modes (**i-iv**), shown for various values of the number of motors on the cargo, *N*. Dashed lines indicating scaling with diameter *d* to the indicated power are shown for comparison. Overlaid curves are fits, detailed in table S1.

We find that 0.5 μm diameter cargos with a single free motor with diffusion coefficient 0.1 μm^2^ s^−1^ bind more than an order of magnitude faster than cargos of the same size with a single rigidly attached motor. This also can be understood by considering timescales: diffusion on the surface is much faster than rotational diffusion of the cargo, so less time is spent waiting for a motor to come near the MT. When free motors are added in a cluster, we find that time to bind does not decrease with more motors in that cluster. In fact time to bind increases slightly. This is due to our assumption that clusters with more motors have a diffusion coefficient that decreases with the number of motors in the cluster (more drag through the cargo surface), see 4.7. This indicates that the time spent waiting for motors to come near the MT is the slowest process.When anchors are independent, time to bind goes down drastically as the number of motors is increased. This effect includes the decreased time to bind from the spread out initial locations of the motors, as well as the fact that each motor performs its own search. While anchor motions are all subject to the same contribution from the rotational diffusion of the cargo, the rotation timescale is much slower than surface diffusion, making each search almost independent at this cargo size and diffusion coefficient.

The results of Fig 3A indicate that at 0.1 μm^2^ s^−1^, surface diffusion is much faster than cargo rotation. At some diffusion coefficient, surface diffusion should become slower than cargo rotation, and the time to bind for free independent cargos should approach that of rigid dispersed cargos. We find that the motor diffusion coefficient must be decreased by orders of magnitude to obtain significant changes in the time to bind, as shown in Fig 3B. For the 0.5 μm diameter cargos shown, diffusion coefficient of the motors must be lower than 10^−4^ μm^2^ s^−1^ for free independent cargos to have times to bind similar to rigid dispersed cargos.

We find that times to bind for cargos in each of the organization modes depend differently on the cargo size. When cargos are small enough that all motors can simultaneously reach the MT (~50nm diameter), the time to bind for cargos in all organization modes is the same. (We note our model assumes no direct, e.g., steric, interaction between motors, which could influence this result strongly for small cargo and many motors.) As cargo size increases, time to bind remains dependent only on motor number until cargos reach ~100 nm in diameter. For cargos larger than this, scaling of time to bind with size is drastically different for cargos in the different organization modes. For cargos with rigidly attached motors, time to bind scales with approximately the fourth power of the cargo diameter. For cargos in the free independent mode, time to bind scales with between the second and third power of the diameter. Free clustered cargos with one motor are identical to free independent cargos with one motor, and have the same scaling. As the number of motors increases, however, scaling becomes more severe, nearing the scaling of rigid motors at high motor number.

### 2.3 Cargos with free independent motors form dynamic clusters which increase travel distance

We next investigate the distance that cargos travel, after initial attachment to the MT. To do so, we begin simulations with a single motor bound to the MT and simulate until the cargo reaches a state in which all motors are detached from the MT. A few stochastic trajectories, along with the mean position over many cargos are shown as a function of time in Fig 4A (top). Once bound to the MT, rigid clustered cargos and free clustered cargos behave similarly. Hereafter, we show only results for rigid clustered cargos and refer to them as “clustered” to reflect this. Visualizations of example simulations for each organization mode are shown in videos 6–8.

**Figure 4:**
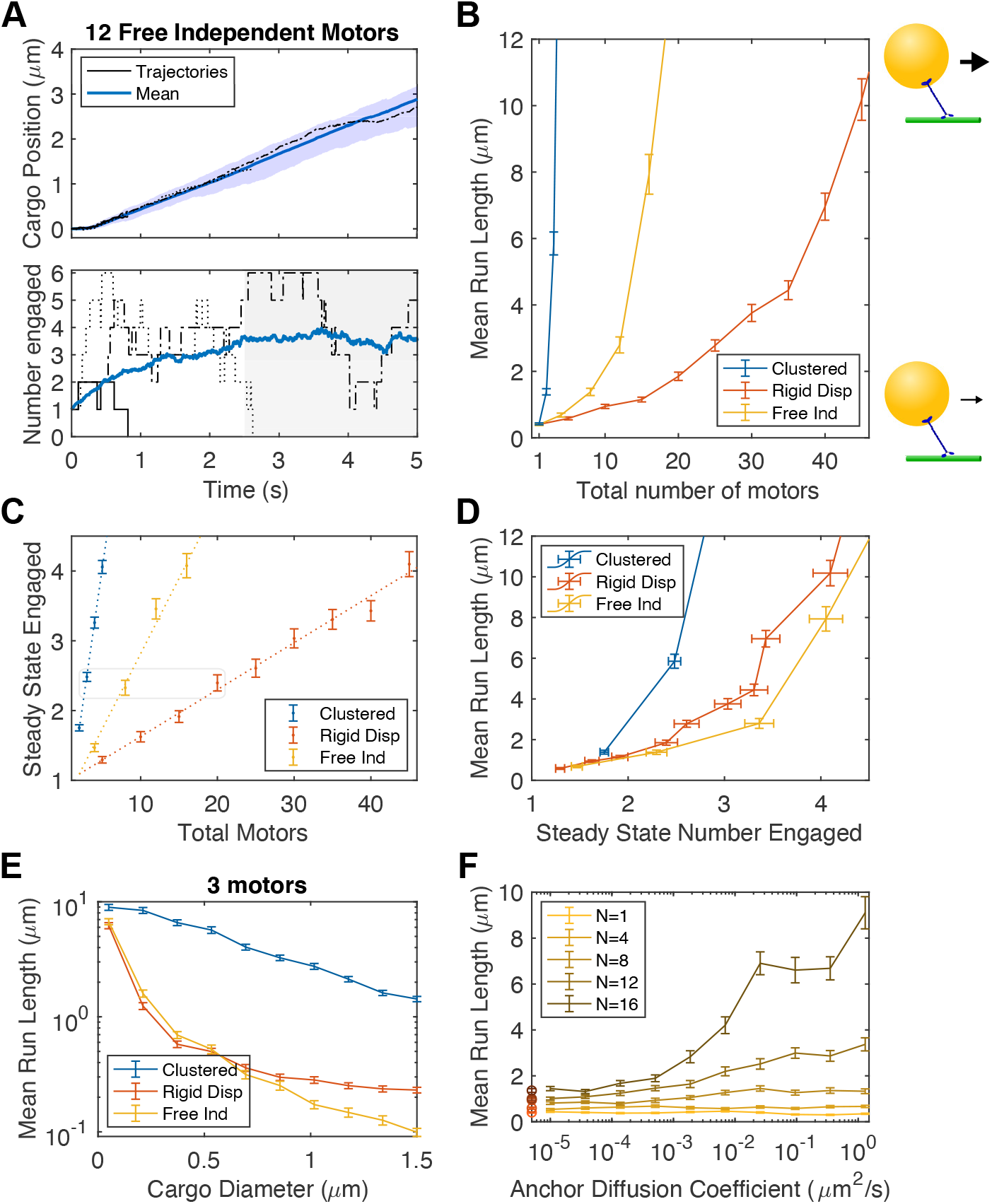
Free independent cargos have longer run lengths than rigid dispersed cargos due to dynamic clustering **A**: Position (top) and number of motors engaged (bottom) with time for free independent cargos with 12 total motors. Trajectories of three cargos are shown in black and the mean over 100 cargos is shown in dark blue. Cargos are excluded from the calculation after they fall off the microtubule. The area between the 5th and 95th percentile positions is shaded in blue in the top panel. Times greater than the time to steady state are shaded in grey in the bottom panel. **B**: Mean run lengths for cargos in each of the three anchor modes as a function of the total number of motors on the cargo. Distributions of run lengths are shown in figure S3A. **C**: Mean number of motors engaged at steady state versus motor number for the 3 anchor modes. Linear fits are shown as dotted lines. For values, see table S1. Grey box indicates conditions with similar mean number engaged at steady state which will be compared in later figures. **D**: Mean run length as a function of steady state number of engaged motors for the three anchoring modes. **E**: Mean run length for cargos with three total motors as a function of cargo size for each of the three anchoring modes. **F**: Mean run length for cargos with free independent motors as a function of the diffusion coefficient of the motors in the cargo membrane, for different numbers of total motors. Red error bars overlaid on the left axis are for rigid dispersed cargos of the same total numbers. In **B, E** and **F**, error bars are SEM of 300 cargos. In **C**, error bars are SEM of 100 cargos. In **B, D, E** and **F** lines between points are guides for the eye.

We find that cargo run lengths depend strongly on motor organization mode. For cargos with clustered motors, just four motors working together give run lengths on the order of the size of a cell. Motors in this mode work together very well, as if any motor is bound the rest of the motors are located where they are also able to bind the MT. This contrasts with the dispersed mode, where many motors are necessary to achieve run lengths of a few microns. For the 0.5 μm diameter cargos plotted in Fig 4B, 25 motors are required to achieve a mean run length of 3 μm. These results are consistent with previous work comparing these two modes [4].

For cargos with free independent motors, we find run lengths which are longer than those of dispersed cargos, but not as long as those of clustered cargos. One possibility is that the run lengths are due to the number of motors which are instantaneously bound to the MT at a given time, which we term the number engaged. We therefore query this quantity in our simulations. The number of motors engaged on the MT fluctuates with time. Several stochastic trajectories are shown in Fig 4A (bottom), along with the mean over 100 cargos at each time. We find that the mean number of motors engaged rises from the initial condition of 1 to a steady state value over a period of time. In Fig 4C, we show that free independent cargos have more motors engaged than rigid dispersed cargos. The initial locations of motors on the surface of cargos in these two modes is the same, i.e., uniformly random on the surface. Therefore, the increased steady state number of engaged motors on free independent cargos indicates that motors are diffusing to the MT, binding, and remaining bound for longer than they would if simply placed randomly. In other words, the motors cluster near the MT. These clusters are dynamic, with motors diffusing in and binding, as well as unbinding and diffusing away, as can be seen in videos 1 and 8.

How strong is the clustering effect? In the range of total motor numbers investigated, the number of engaged motors is 25–30 % of the total number on the cargo. This is more than the 10–15 % of motors engaged on rigid dispersed cargos, but less than the ~ 80 % of motors engaged on clustered cargos (Fig 4C). So, while dynamic clusters contain more motors than would available to bind the MT if motors were distributed randomly on the surface, they do not contain all or even most of the motors on the cargo.

We hypothesized that dynamic clustering is responsible for free independent cargos’ enhanced run length over rigid dispersed cargos. To test this hypothesis, we plot mean run length vs. steady state number engaged. If dynamic clustering is responsible for the enhanced run length, we expect the free independent and rigid clustered modes to have the same run length when the number of engaged motors is the same. We see in Fig 4D that data from the two modes is similar. Run lengths for free independent cargos are in fact slightly lower once corrected for number of engaged motors. This is surprising because cargos with similar mean numbers of motors engaged at steady state have similar distributions of motors engaged, as shown in figure S3B. In supplementary figure S3C, we show that the lower run length is explained by a longer time to steady state and resultantly more cargos which fall off the MT at early times.

The three modes also differ significantly in their dependance on cargo size. In Fig 4E, we show that cargos with clustered motors have a run length that depends only weakly on cargo size, while free independent and rigid dispersed cargos have a more complex dependance.

The run length advantage of free independent cargos over rigid dispersed ones should, like the binding time advantage, be reduced to zero at low anchor diffusion coefficients. In Fig 4F, we show that the diffusion coefficient must be reduced by orders of magnitude to have significant impacts on run length. We find that diffusion coefficients below 10^−4^ μm^2^ s^−1^ are effectively rigid, which is a similar to the threshold we found for time-to-bind.

### 2.4 Cargos with free independent motors are better able to transport against a load compared to cargos with rigid dispersed motors

A cargo’s ability to generate a sustained force is also important for navigating the crowded environment of the cell. In this section, we examine the run lengths of cargos in the different organization modes against a constant force.

As expected, we find that that increasing force decreases the run length of cargos, no matter what the organization mode or number of motors, as can be seen in S4, A. We find that 7 pN of force is sufficient to reduce the run lengths from 20 μm or more to nearly zero in every organization mode.

We now compare the run length of cargos in different organization modes under force. In Fig 5A, we show that cargos with 20 free independent motors have significantly longer run lengths than cargos with the same number of rigid dispersed motors, when subject to forces up to 7 pN. At higher forces, run lengths for these cargos are effectively zero. At this high number of motors, cargos with clustered motors travel long distances, even when loaded with 12 pN or more (Fig 5A, arrows). When the number of total motors is 5, we can see that cargos with clustered motors outperform both rigid dispersed and free independent cargos (S4, B). At this low number of total motors, cargos in rigid dispersed and free independent modes are almost always driven by a single motor, so differences between the two modes are not apparent.

**Figure 5:**
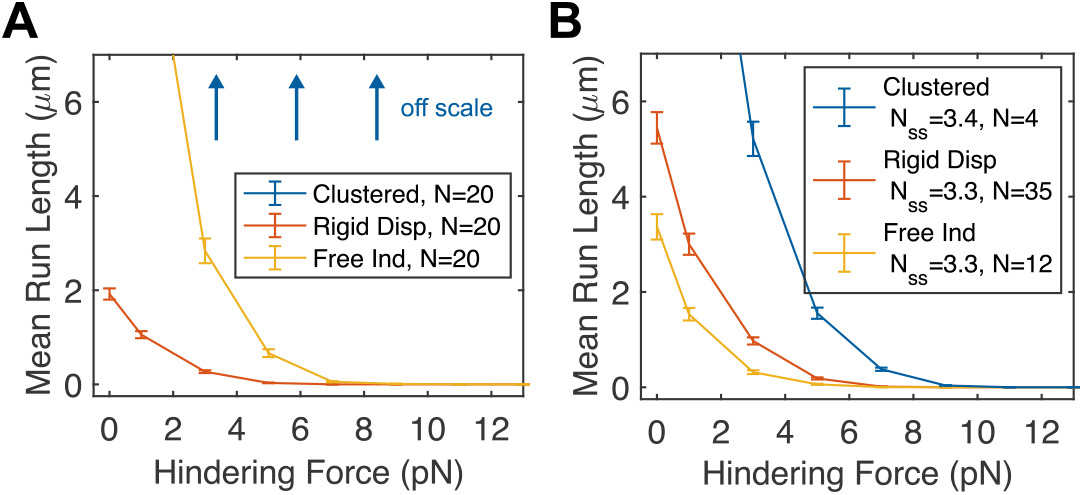
Cargos with free independent motors travel farther under load due to dynamic clustering. Mean run lengths as a function of external hindering load on the cargo for cargos in each of the three modes, with **A**: matched total number of motors. Cargos with 20 clustered motors travel farther than 20 μm on average, and are not shown to clarify the difference between rigid dispersed and free independent modes. **B**: matched number of motors engaged at steady state (at 0 external load). Distributions of run lengths for these cargos can be found in figure S4C. Error bars are standard error of the mean over 300 cargos in both **A** and **B**. Lines between points are guides for the eye.

Is the increase in run lengths under force also solely due to changes in the number engaged? To test this hypothesis, we plot run lengths under force for cargos in the three organization modes with total numbers of motors which give the cargos matching steady state numbers of engaged motors (at 0 force, indicated by box in Fig. 4C). Like in the zero force case, when we compare at the same steady state number, we find that cargos with free independent motors have similar, but slightly lower run lengths than cargos with dispersed motors. Therefore, the enhancement in run length of free independent motors comes from the ability of motors to form dynamic clusters, like in the unloaded case.

A priori, we hypothesized that free motors share the load more equally, and that this would lead to longer run lengths under load, after correcting for number engaged. However, as shown in Fig. 4B, we do not find this effect is strong enough to outperform cargos with rigid dispersed motors on a per-motor-engaged basis.

## 3 Discussion

While transport of subcellular cargo by molecular motors is increasingly understood, the extension of this understanding to control of cell internal organization will involve studying the three-way interplay between the cargo, the MT and local environment, and the motors. In this work, we made a particular effort to simulate behavior in viscosities relevant to the cellular environment. Our work points to parts of cargo transport that are not well understood, and informs the picture of overall transport. We developed a computational model of the motors’ interaction with the cargo, assuming different modes of organization. These modes varied both in the positions of the motors on the surface of the cargo and the freedom of those positions to change via movement in a fluid membrane. Combining Fig. 3 and 4, we arrive at our main conclusion, summarized in Fig. 6. First, cargos with rigidly attached motors face a tradeoff between, on the one hand, time for the cargo to bind to a nearby microtubule and, on the other, the run length of cargos once one motor is bound. Rigid clustered motors bind slowly, but have long run lengths. Rigid dispersed cargos bind faster than rigid clustered cargos, but have short run lengths. Second, depending on parameters, cargos with free motors can overcome this tradeoff. Cargos with free independent motors bind faster than cargos in either of the rigid modes, and have run lengths which are longer than those of rigid dispersed cargos, but not as long as clustered cargos. When motors are arranged in a free cluster, cargos have the same long run lengths as rigid clusters, as well as binding to a nearby microtubule more quickly. The time to bind is not as fast as a free independent cargo, however, or even a rigid dispersed cargo with many motors.

**Figure 6:**
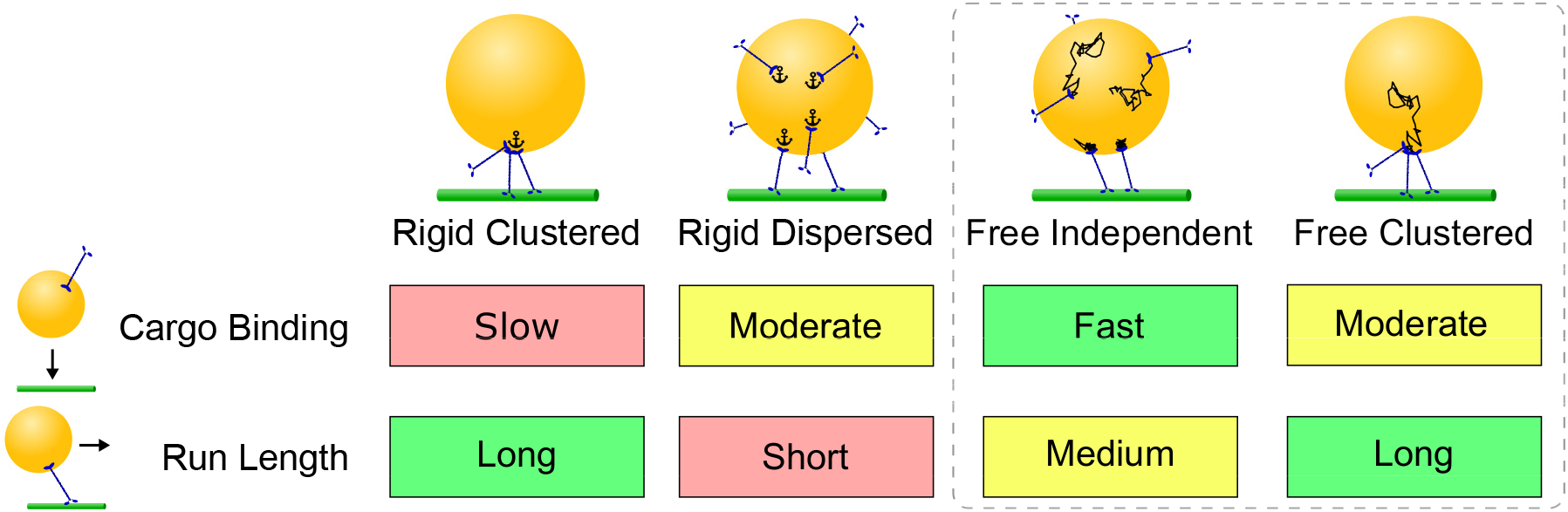
We find that different organizations of the motors on the cargo have different implications for how rapidly the cargo will bind to a nearby microtubule and the cargo’s ability to travel along the microtubule. For cargos with motors rigidly bound, (first two columns) there is a tradeoff between clustered motors, which are slow to bind the microtubule, but travel long distances, and dispersed motors, which bind the microtubule quickly but are have poor travel distances. Cargos with freely diffusing motors (third column), at reasonable surface diffusion coefficients, cargo sizes and motor numbers, overcome this tradeoff. They bind the microtubule at least as fast as rigidly dispersed motors, and faster for large and realistic estimates of diffusion coefficient. They travel farther than rigid dispersed motors because of the formation of dynamic clusters. Because these clusters have high internal turnover, travel distances are lower than that of a rigid cluster. For a cargo with a freely diffusing cluster of motors (fourth column), cargo binding may behave more like a rigid cluster or freely diffusing motors, depending on parameters. Once it has bound the microtubule, it behaves indistinguishably from a rigid cluster in both run length and force generation.

### 3.1 Cargo size provides a potential sorting mechanism and avenue for studying motor-cargo attachment

We find that time to bind and run length are sensitive to cargo size in different ways for the four organization modes (Fig. 3C). Rigid clustered cargos have a time to bind which scales up strongly as cargo size increases, but run length is relatively insensitive to cargo size. The time to bind for rigid dispersed cargos also scales strongly, but their run length is more sensitive to cargo size in comparison to rigid clustered cargos. Free independent cargos have a much weaker scaling of time to bind, as well as an intermediate dependance of run length on cargo size. These different scalings raise the possibility that the cell could use the size dependent behaviors to differentially direct otherwise similar cargos.

Binding time scaling found in Fig. 3C could be used to identify the organization of motors on cargos from the cell. Using the natural variability of cargo sizes to uncover different scaling exponents could inform whether the motors are free or not.

### 3.2 Comparison with previous studies

This work adds to a growing body of evidence that, first, the position of motors on the cargo (whether free or not) impacts transport [4, 24]; second, that mobility on the cargo surface further influences transport [8, 9, 17, 18, 25]; and, third, that there are significant differences in the arrangement of motors on cargo in cells [3, 18], for example via changes to lipid content [26, 27]. The closest direct experimental study related to our work is by the Xu Lab [9]. *In vitro*, they find broad agreement with our simulations that fluid membranes enhance transport. However, the details are different: they find increased velocity for membrane cargos where we predict none, and do not find a run length increase. We note that our simulations predict significantly longer run lengths for free motors only when total motor numbers are more than about 5, a regime not tested in [9]. Our work thus highlights the difficulty extrapolating in vitro results into the cellular environment.

Our work is in broad agreement with Chowdary *et al*. [18], who combined advanced microscopy and tracking with a computational model to give insight into the motion of endosomes in axons. They report the emergence of dynamic clustering in their simulation via preference for binding the MT track. Their simulation allows motors to move on the surface of a cargo independent of each other and of rotations of the cargo itself. Our work extends this by considering a model in which motor anchor position, cargo rotation and cargo translation are all coupled with both cytoplasmic viscosity/fluctuations and membrane viscosity/fluctuations, via a force-balance relationship at every time point. Particularly interesting complexity arises, e.g., the two viscosities give rise to fluctuations that are, in the lab frame of reference, no longer uncorrelated (see supplemental). Doing so allowed us to explore a wide range of parameters, from the limit in which the cargo membrane is nearly inviscid and the motor can step along the MT without significantly moving the cargo, all the way to a static anchoring. It also allowed us to explore the competition between cargo rotation and diffusion of motors to drive binding time – of particular importance when the cargos are small. The coupling via force-balance also led to an emergent inter-motor communication via how each motor is loaded, which influenced predicted run lengths. Interestingly, we find that dynamic clustering is not a strong effect for all parameters regimes. For example, at best guess parameters, we find only 25% of the motors are clustered (e.g., Fig 4).

Our work contrasts with Jiang *et al*. [28], whose model predicts that many motors are necessary to achieve long run lengths. Jiang *et al*. [28] measure the accumulation of diffusing motors on microtubules in a gliding assay configuration, and find that motors bind slowly. In their model, kinesin binds at ~5s^−1^ (which this paper also uses), but has only a short reach. The model for kinesin binding in this paper uses the length of kinesin-1 in electron microscopy for the reach parameter. We consider this to be a lower bound for the maximum possible reach of kinesin in cells, since adaptor proteins likely extend this length somewhat. It is clear that in some situations kinesin binds quickly [5], while in others it binds slowly [28]. It is unclear why, and even more unclear how to model this process in the context of the cytoplasm. Measurements of the number of kinesins on cargos have been made. Photobleaching indicates that just a few kinesins are present on mouse macrophage phagosomes [12] and phagosomes from mouse brain [11]. Quantitative mass spectrometry on synaptic boutons finds that 1-2 kinesins are present per vesicle [29] (see 4.9). While few motors are present on these cargos, it remains unclear how motor numbers vary between cargos of different types and sizes.

We reiterate another subtlety, which is that there are distinct biophysical notions of attachment [5]: The time for a cargo, on first approach to a MT (e.g., from the organelle that generated it), the time to rebind after pulling out of a trap, the concentration-dependent local attachment rates of the motor head to the MT lattice, and the local attachment rate of a second motor to the MT on a multi-motor cargo. It is possible that such subtlety underlies differences between these studies and ours.

### 3.3 Influence of cargo surface mobility on transport

Our results show that changing the diffusion coefficient of motors can change transport. As shown in Fig. 4F, a large change in diffusion coefficient is necessary to significantly change transport (potentially explaining why [28] did not observe a change in their assay, which also had a different geometry that will significantly influence motor behavior). It is tempting to speculate that this could be used to modulate transport. In Sec. 4.6, we discuss theoretical estimates for upper bounds on diffusion coefficients. Diffusion coefficients several orders of magnitude lower have been measured for transmembrane proteins [30], showing that a wide range is achievable in the cell. Briefly, this could be achieved by changes in lipid composition [9, 28], changes in linker molecules [27, 31, 32], membrane tension [33] or by membrane crowding. Whether any of these modulators would lead to a sufficiently large change to modulate transport remains to be determined. However, in a potential analogous system, we note a 5-fold change in membrane diffusion of cdc42 in the yeast outer membrane has been suggested to be under regulation during yeast mating [34]. Another possibility is by changing anchoring to a structure like a lipid raft [3, 27] or cytoskeletal structure [23], which could (if the anchor were long-lived) reduce diffusion coefficient to nearly zero.

### 3.4 Quantitative model provides framework to infer motor organization

It is possible to combine various measurements of cargo and transport properties made in cells with the results of our model to learn about those quantities which are unknown. Data on forces generated in optical traps, which can be used to infer numbers of active motors, is abundant ([6, 13, 35–38], many others). Binding time has been quantified *in vivo* in [6] and *in vitro* in [6, 12]. We are aware of direct reports of (total) motor numbers for two cargos [11, 12], and indirectly for one more ([29], see 4.9). We know of motor organization reported for one case [3]. It is rare that several of these quantities are reported for the same cargos. Chaudhary *et al*. [12][39] report motor numbers, forces, and binding times in *vitro*, from which it may be possible to infer organization. Reddy et *al*. [6] report binding times and forces in cells, which we comment on below. Combining measurements of binding time, force generation, and motor number would allow for significant insights into both how cargos are transported and how kinesin and other motors function in the cell. Since each of these quantities has been measured individually, they need only be combined to allow inference on organization. Combining optical trapping with super-resolution microscopy [40] could allow for even better, individual cargo specific characterization. We point out that super-resolution measurements would likely not be able to fully define organization, since free independent motors may look the same as rigid dispersed motors (or possibly clustered motors) in a snapshot. However, using our results, it would be possible to combine an organization snapshot with force measurements and/or binding times to determine organization.

Both binding times and forces for COS-1 lipid droplets can be obtained from the experiments conducted in [6]. In Sec. 4.4, we revisit data collected in [6] to extract kinesin-driven binding times. The measured 7 s binding time is consistent with either ~5 rigid dispersed motors, or 5 or fewer free independent motors (at varying diffusion coefficients). However, the mean forces generated by the lipid droplets indicate they are often driven by multiple kinesins. We find that for the parameters chosen, both free independent and rigid dispersed cargos with 5 motors are almost always driven by a single motor (Fig. 4C). Extrapolating from the results of our model, two resolutions to this conundrum present themselves. First, if kinesins were organized into groups of 2 which were then rigidly dispersed on the cargo, they would have similar binding times to 5 single motors rigidly dispersed, as binding times are rotation limited. They would also engage 2 motors much more often, resulting in higher forces. Second, free independent motors could form stronger dynamic clusters and engage more motors simultaneously if the binding rate of subsequent motors was higher than 5s^−1^, possibly allowing for both 7 s binding time and multiple motor forces. Since MAP7 is known to increase kinesin binding rate [39, 41, 42], a higher binding rate is possible for the kinesins on the lipid droplets in COS-1 cells.

### 3.5 Future model extensions

That the transport machinery is sensitive to motor organization opens several possible directions of future work. First, cargos in the cell exist in a microtubule network with a specific architecture [16, 43–45]. The current work focused on exploring interaction with a single MT, but could readily be extended. Second, many cargos are deformable [46] and this might lead to significant differences in transport. While deformability of a large cargo will lead to transport changed by interacting with the environment, e.g., spacing of nearby cytoskeletal elements and organelles, our work suggests cargo deformability might also impact transport more directly via changes to motor organization. Finally, motors are sensitive to MT-associated proteins (MAPs) [47, 48]. The accessibility of the motors to MAPs is expected to exhibit similar behavior to the accessibility of motors to the MT, which we have shown to vary widely depending on organization mode.

## 4 Methods

### 4.1 Chemomechanical model of cargo transport with viscous membranes

In [16], we presented a chemomechanical model for cargo transported by multiple motors. The cargo is modeled as a sphere in a Newtonian fluid with cytoplasmic viscosity, that can translate and rotate under deterministic forces and thermal (stochastic) forces. Motors bind, unbind and step in stochastic, force-dependent and location-dependent manner.

Here, to simulate motors diffusing on the surface of the cargo, we extend the model from [16] in the following ways. First, we allow the motor anchor points to move on the surface of the cargo, experiencing a second viscosity in the local plane of the spherical cargo. This increases the number of degrees of freedom in the computation from 6 (translation and rotation of the cargo) to 6 + 2*N* where *N* is the number of motors (in addition to the location of the motor heads). To describe motor diffusion, we also must now consider stochastic thermal forces in the second viscosity. Finally, we also develop new force-dependence for stepping and unbinding in response to experimental evidence, e.g., [19, 20].

To numerically simulate this new model, in which motors experience two viscosities and two distinct modes of thermal fluctuations, we first formalize the equations of motion (not including binding, unbinding and stepping) and solve these analytically, then input these into the numerical scheme that uses an Euler-Maruyama time-stepping for motion and a Gillespie-type scheme for binding, unbinding and stepping.

Full description of the model is in the Supplemental Material, and the code is openly available at github.com/mbovyn/Motor-Cargo-Simulator (DOI:10.5281/zenodo.4325111).

### 4.2 Estimate for viscosity experienced by cargos rotating in the cytoplasm

The cytoplasm is known to have complex rheology, with estimates for its viscosity varying orders of magnitude [49]. In Supporting Material section A2.1.3, we provide an in-depth literature review and discussion of estimates for cytoplasmic viscosity. We conclude that the most relevant study is Wilhelm *et al*. [50], who used active, magnetic rotational microrheology to study chains of magnetic endosomes in HeLa cells. The authors report a value for the viscous component of the resistance to rotational motion as 2 Pa s. Since the chains of endosomes are several microns long and are possibly experiencing resistance from being dragged through the cytoskeleton, it is unclear if spherical cargos would experience similar resistance. Wilhelm et al. then repeat their measurement on cells with cytoskeletons perturbed by either latrunculin A or nocodazole, finding that in either case the viscous component is reduced to 0.4 Pa s. We select this value as our best guess for the viscosity experienced by rotating cargos. For further discussion, see Supporting Material section A2.1.3.

### 4.3 Extrapolation of rebinding time from previous work *in vitro*

In [5], we measure the time it takes for beads with single kinesin motors to bind to the microtubule after detaching in an optical trap. We find the time increases with viscosity, from 0.8 s in water to 4. 4 s at 10x water viscosity (0.01 Pa s). We construct and fit two models of the binding process, one assuming motor binding is reaction limited and the other assuming motor binding is diffusion limited. When we evaluate these two models to 0.4 Pa s, we find times of 100 s and 300 s, respectively. We additionally fit a line to rebinding time as a function of viscosity and extrapolate to 0.4 Pa s to find 190 s.

### 4.4 Quantification of rebinding time for lipid droplets

We use data from experiments originally conducted in [6], which focused on dynein-driven cargo, to estimate kinesin-associated binding times. Forces produced by the motors on the lipid droplets (LDs) from COS1 cells was measured using an optical trap (700 mW, 980 nm single mode diode laser, EM4) and Position Sensitive Diode(PSD, First Sensor CA) assembled onto an high resolution DIC Nikon microscope, as described in [6]. High resolution real time particle tracking and automated translation stage were used to stall a linearly moving LD inside the cell by positioning the optical trap on to its center. The motors on the trapped LDs produce a series of stalls and detachments from MTs in the cell. MT orientation in the COS1 cells is majority plus ends out. So whenever the LD moved linearly outward from the cell center they are predominantly moved by kinesin motors (For quantification they are designated as MT Plus end or P events) and dynein motors move the LDs towards the cell center (MT Minus end or M events).

In COS1 cells motor detachment and rebinding events can be observed as clear peaks and sharp step like falling of PSD voltage signal in the PSD (2 kHz). Particle tracking of LDs in DIC video (30 Hz) also showed similar position displacements. A trap stiffness of 5–8pN/100nm and cell thickness of ~1–3 μm are found to be optimum conditions for force measurements. For *in vitro* force measurements, LD motion was reconstituted after purification from COS1 cells using a five step sucrose gradient centrifugation [6].

In table 1, we report summary statistics for the measured binding times. PP represents the average time between LD plus end detachment and resumption of its motility in plus end. MP is the average time between LD minus end detachment and resumption of its plus end motility inside the optical trap.

**Table 1:**
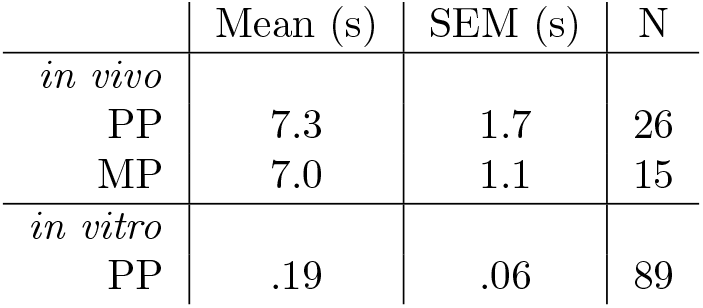
Measured rebinding times for lipid droplets in COS1 cells and for lipid droplets purified from COS1 cells and with their motility reconstituted *in vitro*. Mean and standard error of the mean (SEM) taken from N binding events. Quantification was done on data taken for [6].

|  | Mean (s) | SEM (s) | N |
| --- | --- | --- | --- |
| *in vivo* |  |  |  |
| PP | 7.3 | 1.7 | 26 |
| MP | 7.0 | 1.1 | 15 |
| *in vitro* |  |  |  |
| PP | .19 | .06 | 89 |

### 4.5 Time for a motor to reach the microtubule

In figure 1 we show an estimate of the time it would take for a motor, initially opposite the MT, to come within reach of it. These times come from analytical expressions for mean first passage time to a spherical cap,

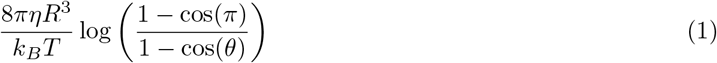

for rotational diffusion of the cargo and

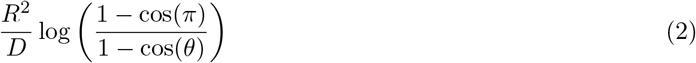

for diffusion in the cargo surface [51]. Here *η* represents viscosity (of the fluid surrounding the cargo), *R* is the cargo radius, *k_B_* is Boltzmann’s constant, *T* is temperature, and *D* is diffusion coefficient (of the motor in the cargo surface). To apply these equations, we had to first estimate *θ*, the extent of the spherical cap to which the motor must diffuse before binding. To do so, we simulated motors diffusing from the north pole of the cargo and recorded the first anchor location where the motor was able to bind with its maximum rate as described in supplemental section A1.3.3, referred to as a max reach location. We then used the mean elevation of these points as our estimate for *θ*.

Max reach locations and mean elevations are shown for various cargo sizes in figure S1.

### 4.6 Range of accessible motor diffusion coefficients

As discussed in Introduction and Discussion, the diffusion coefficient of motors in the cargo surface is unknown. In Discussion, we mention molecular phenomena that could influence motor mobility.

Here, we perform a reductionist estimate. Saffman and Delbrück described that the diffusion coefficient of a disk in a membrane depends on the size of the disk, the viscosity of the membrane and the viscosity of the surrounding fluid [52]. The highest achievable diffusion coefficient for any membrane viscosity is *k_B_T*/(16*aη*), where *a* is the radius of the disk and *η* is the is the viscosity of the surrounding fluid. The lowest achievable diffusion coefficient approaches 0 as membrane viscosity increases (see [53, 54]). Because both the motor anchor and membrane viscosity are likely highly variable for different cargos, we annotate figure 1B with an estimate for the highest achievable diffusion coefficient (1.3 μm^2^ s^−1^ for *a* = 0.5 nm), and the lowest possible (0 μm^2^ s^−1^).

We explore this full range of surface mobilities in the Results.

### 4.7 Model for free clusters

Keeping with the Saffman and Delbrück model [52], we model a free cluster as having motors grouped together diffusing in a membrane with the same viscosity as motors in the free independent mode. A diffusion coefficient of 0. 1 μm^2^ s^−1^ with a hydrodynamic radius of 1 nm corresponds to a membrane viscosity of 0.009 Pa s μm with the closed form expression of [54]. We are then left to model how the hydrodynamic radius changes as a function of the motor number. We model each new motor in the cluster as adding an additional 2 nm disk to the cluster. We then model the arrangement of these disks as the densest packing possible and take the hydrodynamic radius of the cluster to be the radius of the smallest circle which can circumscribe these disks, given in [55]. For 20 motors, this gives a hydrodynamics radius of 5 nm. Since the resulting spread is small compared to the length of the motor’s reach (80 nm), we approximate all the motors as overlayed at the same point in space. To simulate this, we give a single motor a binding rate *N* times the base binding rate.

Since neither linkers nor clusters are well understood [27], there is no model for cluster arrangement suggested by data. While an individual linker could be bound to more than one motor, reducing this size, it is unlikely a single linker could effectively transmit large forces created by many motors. Packing could also be less dense than ideal, increasing the hydrodynamic radius.

### 4.8 Curve fitting

All fits were accomplished with the fit function in Matlab.

Data in figure 3A were fit to the functional forms in table S1.

Data in figure 3C exhibit two regimes, for small cargos binding time is insensitive to cargo size, while depending strongly on cargo size for bigger cargos. To reflect this, two curves are represented. For small diameters, 1/(N*k*^on^) is plotted. Power laws are fit to all cargos above a certain size for each motor number, and the two curves are joined where they intersect. Power law fits were made to cargos above 50 nm, for rigid clustered and free clustered modes, while fits were made to cargos above 200 nm diameter for rigid dispersed and free independent modes. Power law exponents are given in table S1.

### 4.9 Inference on numbers of motors from published data

Wilhelm *et al*. [29] use quantitative mass spectrometry to report numbers of proteins in synaptic boutons. Present in this data are measurements for kinesin proteins. To estimate the number of motors present per vesicle, we used the instructions included in table S3 to combine their measurements with molecular masses (from UniProt) to find numbers of proteins per bouton. We then divided this total number by the number of vesicles per bouton reported in the paper. Kif5b is present at .1/vesicle, while the most abundant is Kif2a, a 1/vesicle. Kif2a is a kinesin-13 protein reported to transport neuronal vesicles [56].

## 5 Acknowledgments

This work was supported by NIH R01 GM123068 to JA and SG, NIH T32 Training Grant EB009418-07 to Arthur Lander and Qing Nie, the UCI Center for Complex Biological Systems, the BEST IGERT program funded by the National Science Foundation DGE-1144901, NSF grant DMS 1763272 and a grant from the Simons Foundation (594598, QN).

---

**Figure S1:**
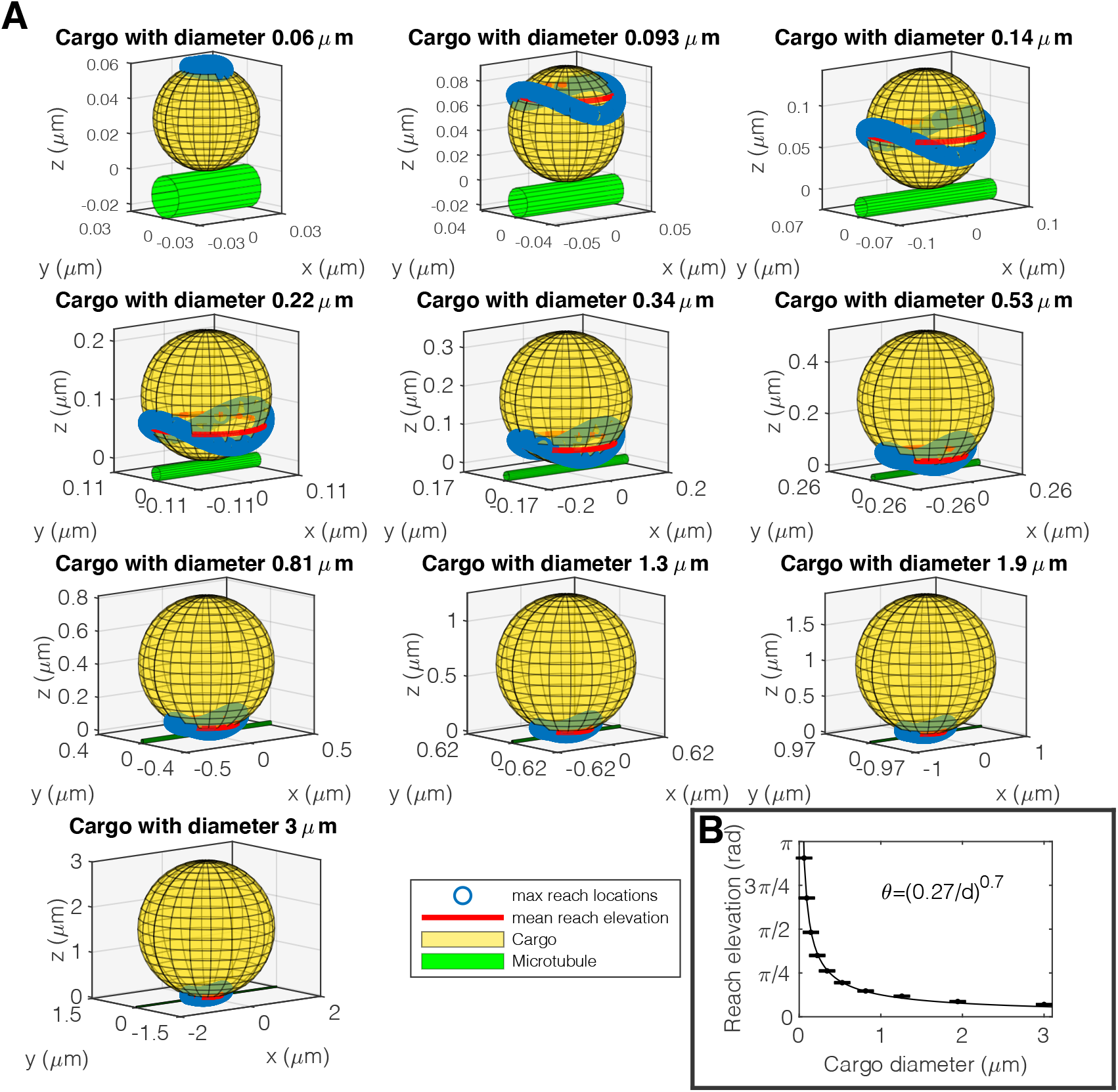
Mean reach elevation, the approximate location on the cargo that a motor can be bound while able to reach the microtubule, used in calculations shown in Fig. 1. Cargo and microtubule were situated as shown in **A**. For details on how we model the mechanics of a motor and its attachment to the cargo, see supplemental section A1.1.1. **A**: Mean reach elevation for cargos of various sizes, as labeled in each panel. Legend in final panel applies to every other panel. **B**: Mean reach elevation for each cargo size shown in **A**. Bars represent standard error of the mean. Data are well fit to reach elevation *θ* = (.27/*d*)^.7^, where *d* is cargo diameter (black curve).

**Figure S2:**
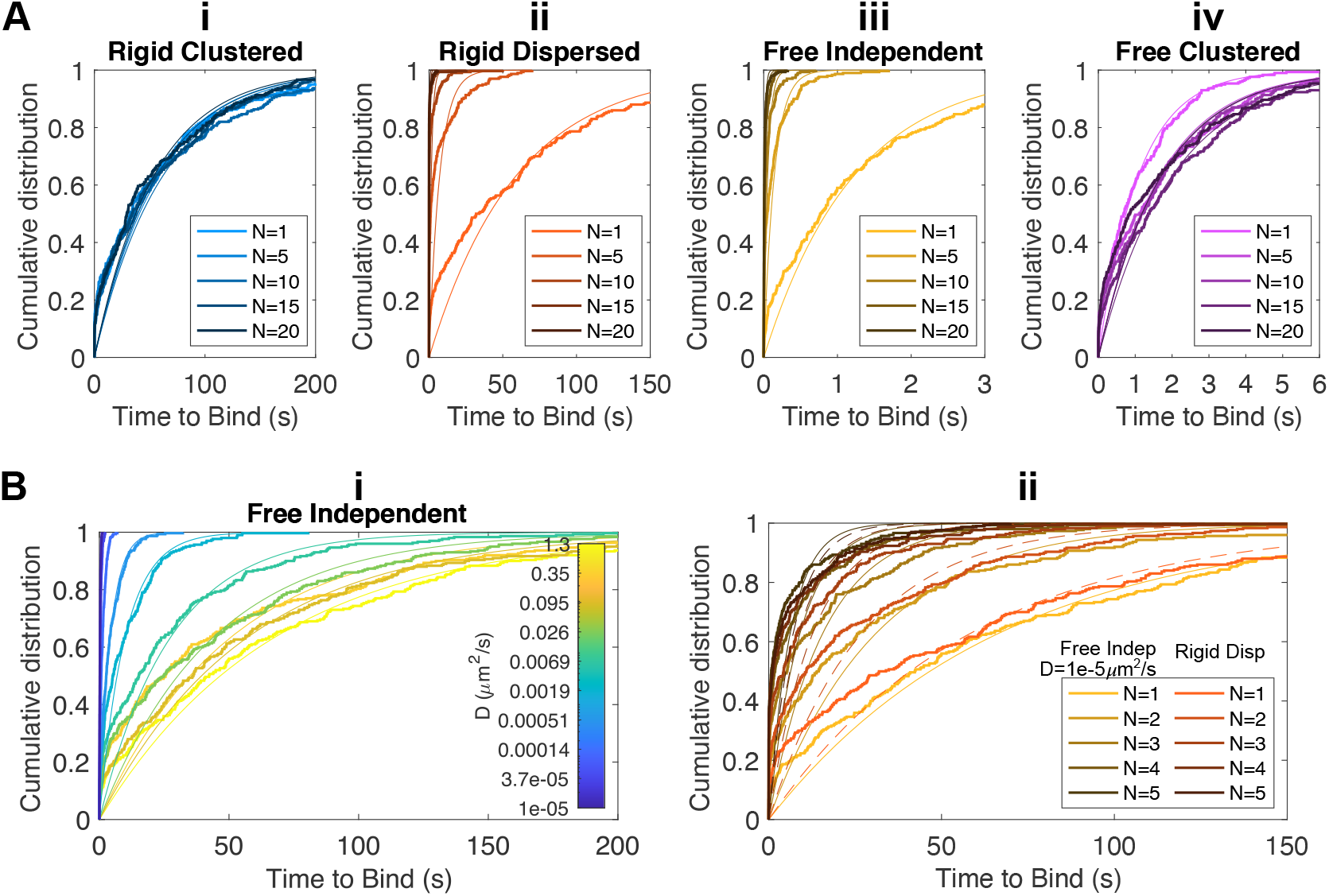
Distributions of times for cargos to bind to the microtubule **A**i-iv: Simulated empirical cumulative distributions of time to bind for cargos in each of the four organization modes. Thinner curves of the same color are exponential fits to the data. **B**i: Simulated empirical cumulative distributions for free independent cargos with a range of motor diffusion coefficients *D*. Diffusion coefficients are log spaced, values of *D* for each distribution shown are labeled on the colorbar. Thinner curves of the same color are exponential fits to the data. **B**ii: Simulated empirical cumulative distributions of times to bind for free independent cargos at diffusion coefficient *D* =1 × 10^−5^ μm^2^ s^−1^ for several numbers of motors on the cargo (yellows). Also shown are distributions for rigid dispersed cargos at the same numbers of total motors (reds). Thinner curves of the same color are exponential fits to the data. Dashed curves are fits for rigid dispersed cargos.

**Figure S3:**
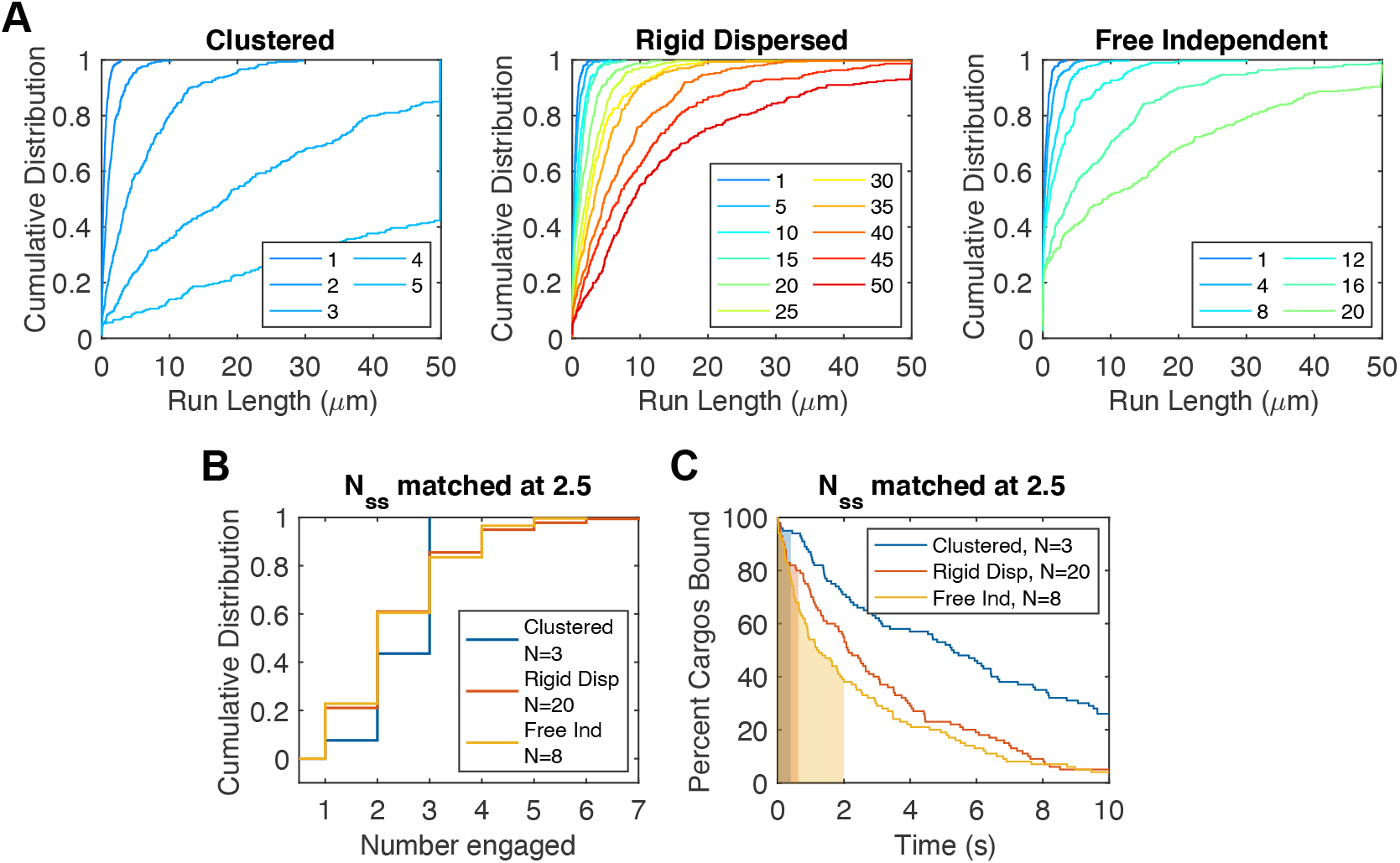
Distributions of run lengths **A**: Simulated empirical cumulative distribution of run lengths for cargos in the clustered, rigid dispersed and free independent modes. Different total numbers of motors are shown on a common color axis. **B**: Simulated empirical cumulative distributions of cargos with closely matched mean numbers of motors engaged at steady state (*N_ss_*). Total numbers of motors *N* was picked for each mode to match an *N_ss_* of 2.5. **C**: Percent of cargos bound as a function of time for cargos with the same total numbers of total motors N as in **B**, picked to match mean number of motors engaged at steady state, *N_ss_*. The time before steady state is reached is filled in color below the curve for each organization mode.

**Figure S4:**
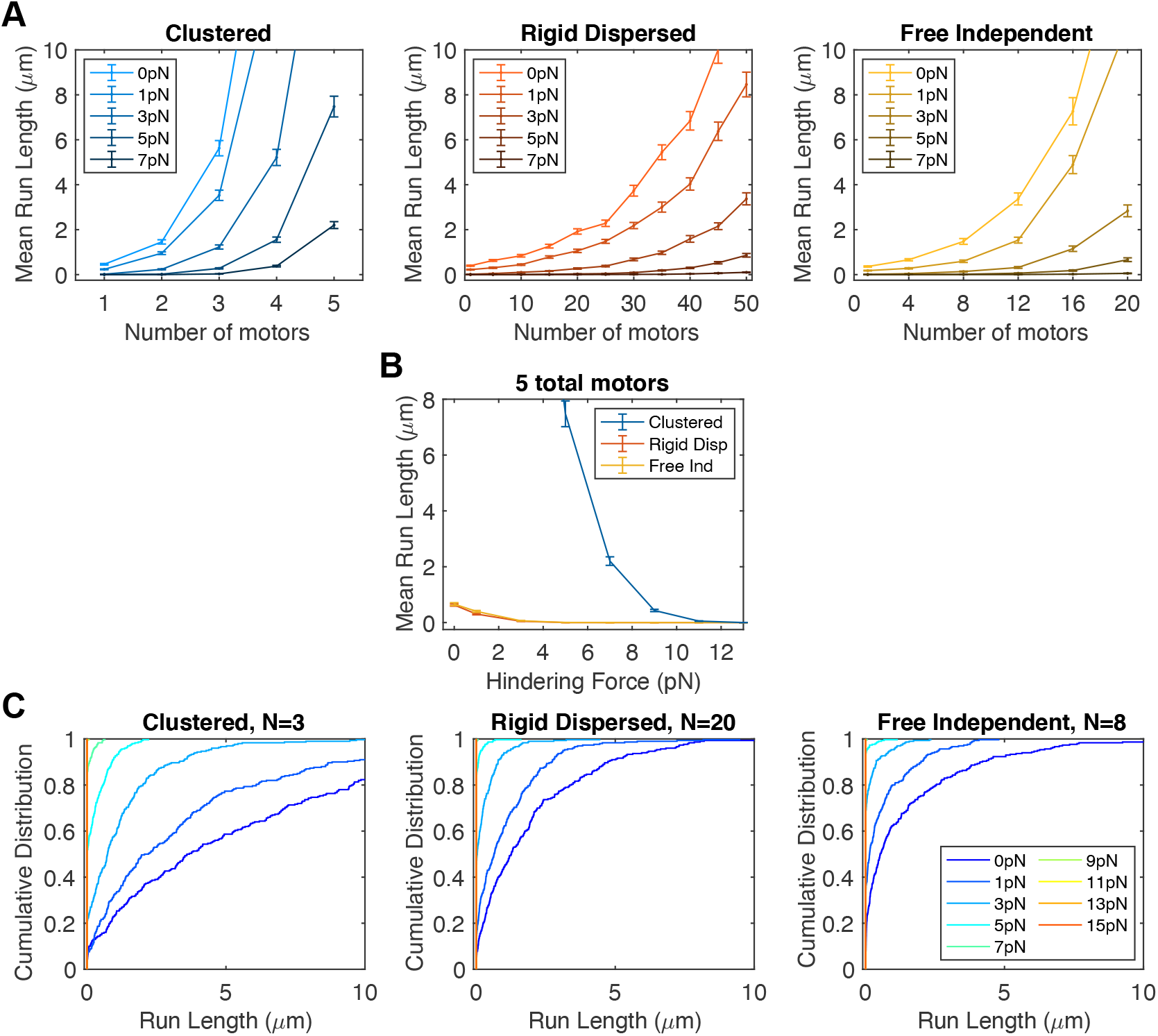
Means and Distributions of run lengths under force **A**: Run lengths as a function of the number of total numbers on the cargo for several values of hindering force on the cargo. Error bars are standard error of the mean. **B**: Mean run lengths as a function of external hindering load on the cargo for cargos in each of the three modes, with the total number of motors on the cargo matched at 4. **C**: Simulated empirical cumulative distributions of run lengths of cargos under different loads. Numbers of motors are chosen to match figure 5B. Distributions are colored on a common axis corresponding to the legend in the right (free independent) panel.

**Table S1:**
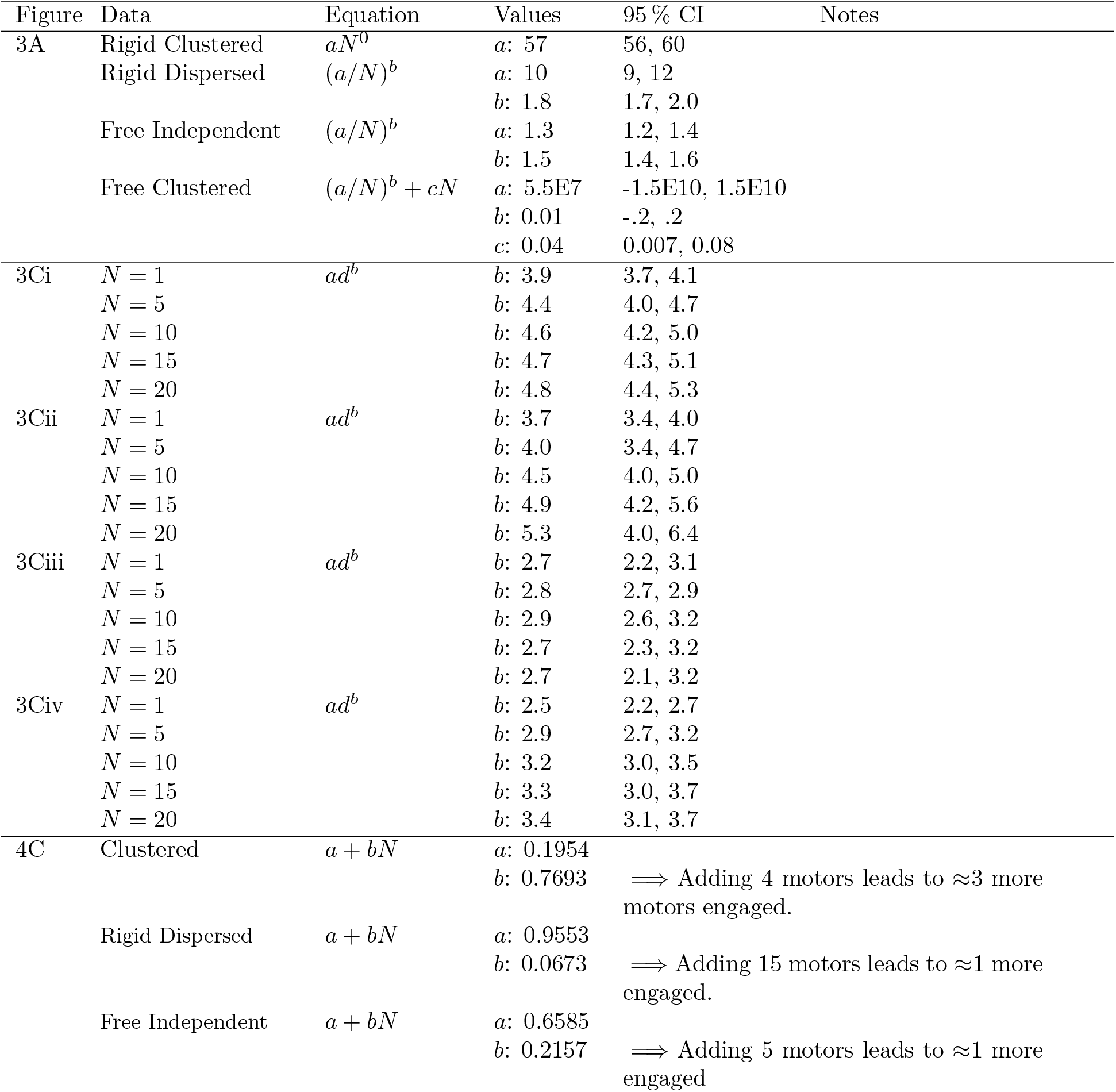
List of fit values

| Figure | Data | Equation | Values | 95 % CI | Notes |
| --- | --- | --- | --- | --- | --- |
| 3A | Rigid Clustered | $aN^0$ | $a$: 57 | 56, 60 | |
| | Rigid Dispersed | $(a/N)^b$ | $a$: 10 | 9, 12 | |
| | | | $b$: 1.8 | 1.7, 2.0 | |
| | Free Independent | $(a/N)^b$ | $a$: 1.3 | 1.2, 1.4 | |
| | | | $b$: 1.5 | 1.4, 1.6 | |
| | Free Clustered | $(a/N)^b + cN$ | $a$: 5.5E7<br>$b$: 0.01<br>$c$: 0.04 | -1.5E10, 1.5E10<br>-.2, .2<br>0.007, 0.08 | |
| 3Ci | $N = 1$ | $ad^b$ | $b$: 3.9 | 3.7, 4.1 | |
| | $N = 5$ | | $b$: 4.4 | 4.0, 4.7 | |
| | $N = 10$ | | $b$: 4.6 | 4.2, 5.0 | |
| | $N = 15$ | | $b$: 4.7 | 4.3, 5.1 | |
| | $N = 20$ | | $b$: 4.8 | 4.4, 5.3 | |
| 3Cii | $N = 1$ | $ad^b$ | $b$: 3.7 | 3.4, 4.0 | |
| | $N = 5$ | | $b$: 4.0 | 3.4, 4.7 | |
| | $N = 10$ | | $b$: 4.5 | 4.0, 5.0 | |
| | $N = 15$ | | $b$: 4.9 | 4.2, 5.6 | |
| | $N = 20$ | | $b$: 5.3 | 4.0, 6.4 | |
| 3Ciii | $N = 1$ | $ad^b$ | $b$: 2.7 | 2.2, 3.1 | |
| | $N = 5$ | | $b$: 2.8 | 2.7, 2.9 | |
| | $N = 10$ | | $b$: 2.9 | 2.6, 3.2 | |
| | $N = 15$ | | $b$: 2.7 | 2.3, 3.2 | |
| | $N = 20$ | | $b$: 2.7 | 2.1, 3.2 | |
| 3Civ | $N = 1$ | $ad^b$ | $b$: 2.5 | 2.2, 2.7 | |
| | $N = 5$ | | $b$: 2.9 | 2.7, 3.2 | |
| | $N = 10$ | | $b$: 3.2 | 3.0, 3.5 | |
| | $N = 15$ | | $b$: 3.3 | 3.0, 3.7 | |
| | $N = 20$ | | $b$: 3.4 | 3.1, 3.7 | |
| 4C | Clustered | $a + bN$ | $a$: 0.1954<br>$b$: 0.7693 | ⇒ Adding 4 motors leads to $\approx 3$ more motors engaged. | |
| | Rigid Dispersed | $a + bN$ | $a$: 0.9553<br>$b$: 0.0673 | ⇒ Adding 15 motors leads to $\approx 1$ more engaged. | |
| | Free Independent | $a + bN$ | $a$: 0.6585<br>$b$: 0.2157 | ⇒ Adding 5 motors leads to $\approx 1$ more engaged | |

## Supplemental Video Descriptions

Cargo, motors and microtubule represented as in 2. Cargo is shown slightly transparent to allow visualization of all motor locations. Hemisphere of motor reach located inside the cargo is an artifact of visualization with transparent cargo.

**S1 Video. Animation from example simulation of cargo with free independent motors.** Frames in Fig 2B taken from this simulation.

### Time to bind

Representative trajectories for time to bind simulations. All times to bind are shown with the same timestep per frame to give context to the differences between modes.

**S2 Video. Animation of cargo binding for cargo with rigid clustered motors.** Cargo eventually binds at *t* =36 s, full trajectory not shown.

**S3 Video. Animation of cargo binding for cargo with rigid dispersed motors.**

**S4 Video. Animation of cargo binding for cargo with free independent motors.**

**S5 Video. Animation of cargo binding for cargo with free clustered motors.**

### Run length

Representative trajectories for run length simulations.

**S6 Video. Animation of run length for cargo with rigid clustered motors.**

**S7 Video. Animation of run length for cargo with rigid dispersed motors.**

**S8 Video. Animation of run length for cargo with free independent motors.**

## A1 Mathematical Model of Cargo Transport with Motor Freedom

**Figure A1:**
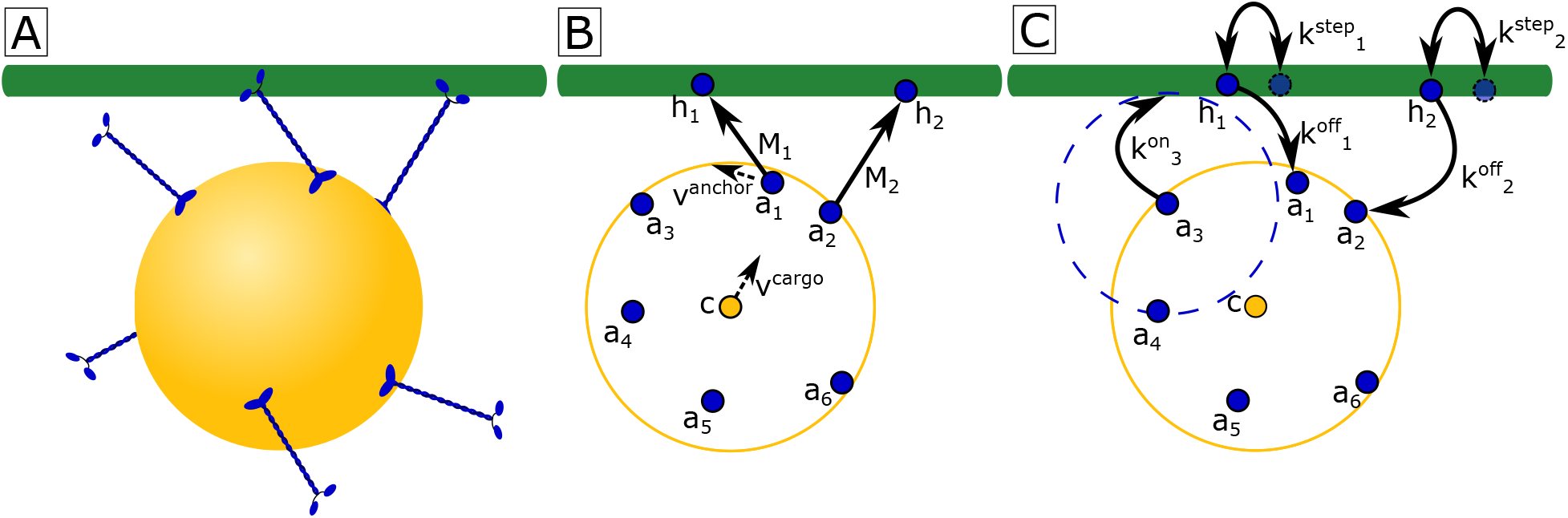
Cartoon diagramming various aspects of the model **A:** We model the cargo as a 3D sphere which diffuses rotationally and translationally. Attached to the cargo are a number of motors which are free to diffuse along the surface of the cargo. **B:** Motors which are bound to the MT exert forces as tension-only springs with a rest length equal to the length of the motor. These forces act to drag the anchors through the membrane, as well as to translate and rotate the cargo. **C:** Motor which are bound to the MT have rates of stepping and unbinding that depend on the force experienced by the motor. Unbound motors have a non-zero binding rate only if the MT is within the motor’s reach.

We present a three dimensional mesoscale model for the dynamics of the cargo and motors. The model takes into account translational diffusion of motors in the cargo membrane, rotational and translational diffusion of the cargo body, as well as stochastic stepping and binding/unbinding of the motors. The locations of the cargo and the anchors where motors are attached to it are governed by a set of stochastic ordinary differential equations. Binding state and location of the motor head along the microtubule (MT) are considered discrete states and transitions between them occur stochastically, modeled as Poisson processes. A cartoon showing general aspects of the model is shown in figure A1. We construct a Monte Carlo numerical simulator, based on a hybrid Euler-Maruyama-Gillespie scheme, and simulate an ensemble of stochastic trajectories, from which we derive transport statistics.

Motors in the model have two states:

Bound: A motor *i* is defined by two points: one represents the motor domains, which we call the head 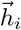; the other represents the location at which the motor is attached to the cargo, which we call the anchor 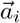. The motor has a rate of stepping along the MT 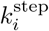 and a rate of unbinding from the MT 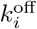.
Unbound: A motor *i* is defined by one point, the anchor location 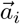. The motor has a rate of binding to the MT, 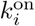, that depends on its location relative to the MT as described below.

We track the anchor locations 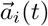, head locations of bound motors 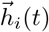, the location of the cargo center 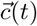, and the cargo orientation 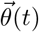. We note that these quantities are time dependent and suppress the (*t*) below where appropriate to simplify notation.

### A1.1 Stochastic ordinary differential equations for cargo motion

We first impose force-balance on the cargo and anchors. The motion of the cargo is in the inertialess regime (Reynolds number ~ 10^−6^), so we omit the inertia term. Similarly, we assume anchor motion is dominated by linear drag and also in the inertialess regime. All forces must balance, so for each anchor

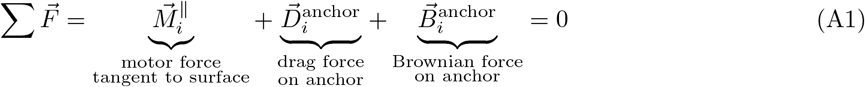

For the cargo body,

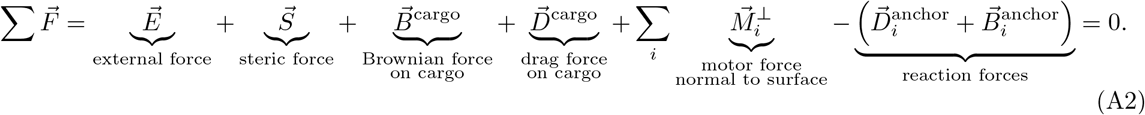

Similarly, we can write the balance of torques on the cargo

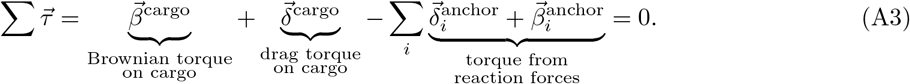

By specifying the forms of each of these forces below, we construct the equations of motion of the cargo center 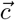, cargo orientation 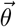, and anchor locations 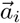.

#### A1.1.1 Motor mechanics and forces

We model the attachment at the motor anchor as a free hinge, in both rotation and bending. Therefore we calculate the distance from the anchor to the MT as the shortest distance which does not imply the stalk would pass though the cargo (see equation A28). We do not include any torque at the anchor.

In stretching, we model the force that the motor exerts as originating from the stretch in spring-like motor stalks, based on experimental measurements [57-59] and in line with previous models [4, 15, 60]. The form of this force is that of a tension-only spring with stiffness *κ*^motor^ and rest length *L*. The force 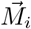 exerted by motor *i* is given by

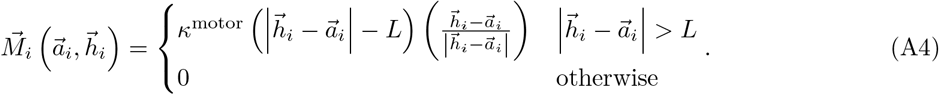

We then break the motor force into components normal and tangent to the sphere at the point of the anchor. We model the cargo as undeformable, so the normal component of the motor force, 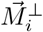, acts to translate the cargo body. The tangential component, 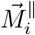, acts to drag the anchor through the cargo membrane. We calculate these components as

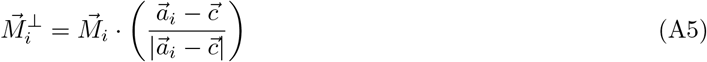

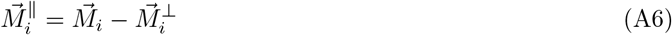

#### A1.1.2 Steric forces

We model the MT as an infinite cylinder of radius *R*_MT_. The cylinder location is defined by a point 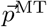 and direction vector 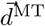. The cargo and MT are prevented from overlapping in space by a steric force with the form of a compression-only spring of stiffness *κ*^steric^ and rest length 0, given by

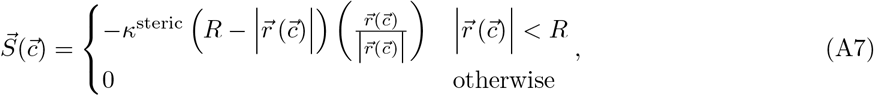

where 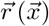 is a vector from point 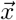 to the nearest point on the MT. This is found by computing the perpendicular distance from 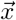 to the MT surface, given by

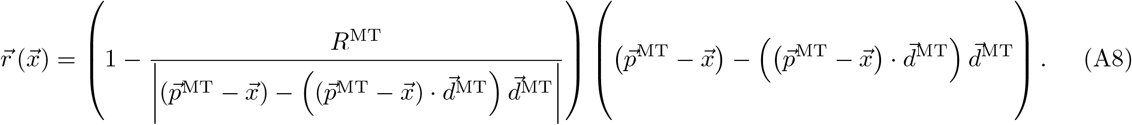

#### A1.1.3 Drag forces

We model the fluid surrounding the cargo as a Newtonian fluid with viscosity *η*. The drag on the cargo is thus given by Stokes’ Law,

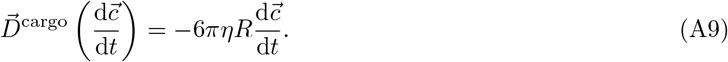

The viscous torque is given by the rotational analogue to Stokes’ Law,

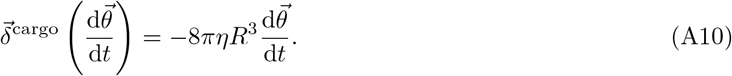

The motion of the anchor in the membrane is also assumed to be dominated by linear drag. The drag force is proportional to the difference in velocity between the anchor and the fluid that makes up the cargo membrane, given by

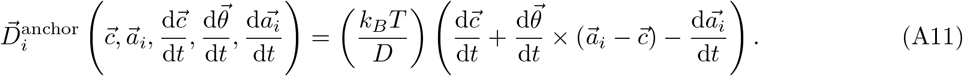

This leads to a torque on the cargo given by the cross product of the lever arm and the force,

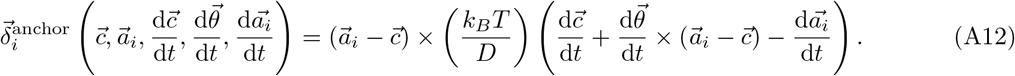

#### A1.1.4 Brownian forces

We model the fluid surrounding the cargo as a Newtonian fluid with viscosity *η*. The Brownian force on the cargo, 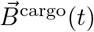, is a vector of three random variables in time (one per spatial dimension) with mean 0 and variance *2k_B_Tξ*, where *ξ* is the drag coefficient and *k_B_T* is the thermal energy unit. Specifically, for the Brownian force on the cargo body,

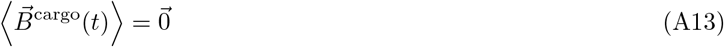

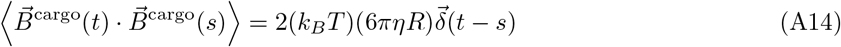

where we have inserted the translational drag coefficient of a sphere at low Reynold’s number and 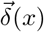 is a vector of three Dirac delta functions. Similarly, the Brownian torque on the cargo, 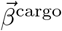, is characterized by

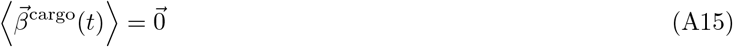

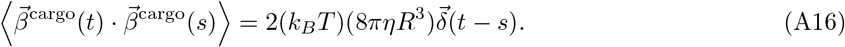

where we have inserted the rotational drag on a sphere at low Reynold’s number.

We model the membrane of the cargo as a Newtonian fluid, in which motor anchor diffuse with coefficient *D*. The Brownian force on anchor *i* in the membrane is characterized by

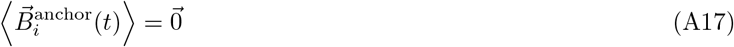

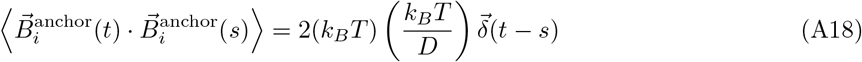

where we have expressed the drag coefficient as the ratio of the thermal energy unit and the diffusion coefficient of the motor in the membrane via the Einstein relation. This force causes a torque on the cargo equal to the cross product of the lever arm and the force,

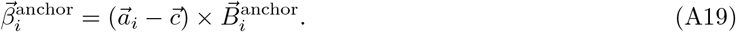

### A1.2 Construction and discretization of the stochastic ordinary differential equations

With the forms of forces specified, equations A1, A2, and A3 represent a set of 3(*N* + 2) coupled ordinary stochastic differential equations, where *N* is the total number of motors on the cargo. The forms of these equations are, for each motor,

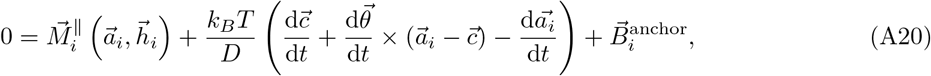

for translation of the cargo,

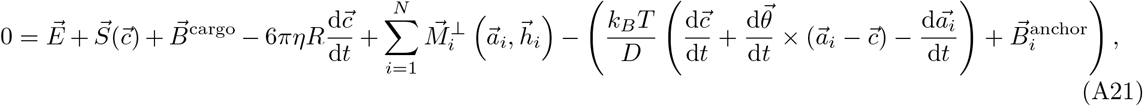

and for rotation of the cargo

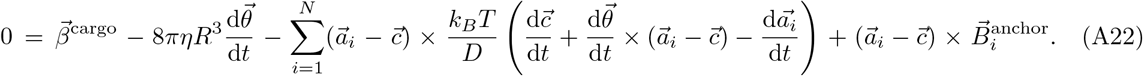

Equations A20 and A21 are specific implementations of the overdamped Langevin equation, used in Brownian dynamics. Equation A22 is the rotational counterpart of A21. The dynamic variables of these equations are the 3*N* anchor position components, the 3 cargo position components, and the 3 cargo orientation components. The system of stochastic ordinary differential equations is linear, allowing us to solve for the derivative terms. The resulting equations have many terms and are not amenable to display.

We discretize these equations according to the Euler-Maruyama method. For an update from the *n*th timestep at time *t_n_* to the next time *t*_*n*+1_ with Δ*t ≡ t*_*n*+1_ − *t_n_*, the discretization is

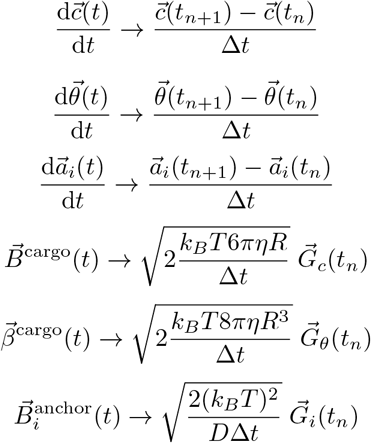

where 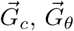 and the 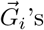 (of which there are *N*) are mutually uncorrelated vectors of three independently and identically distributed gaussian random variables with mean 0 and variance 1.

We note that the update step moves anchor locations in a direction tangent to the surface of the sphere at point *a_i_*(*t_n_*). The new anchor location, *a_i_*(*t*_*n*+1_), then lies outside the surface of the sphere. We force the anchor location back onto the surface of the sphere by correcting it to

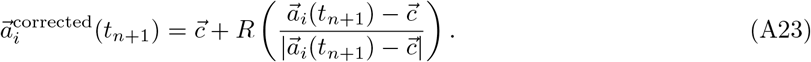

This correction underestimates the true distance the anchor should travel over the surface of the sphere. The magnitude of this underestimation is small for moves where 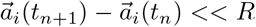. To ensure this is true in simulation, we limit anchor movement distance by choosing a small timestep. To ensure the timestep is small enough, we force the simulation to exit in an error if 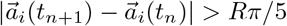 (corresponding to an underestimate of 10%), or if 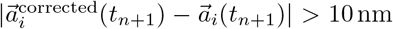. Furthermore, we demonstrate in section A3 that the simulation shows good agreement with analytical calculations of mean first passage times for diffusion on a sphere.

### A1.3 Poisson processes

We model all state transitions in the system as Poisson processes. Experiments have reported exponential distributions of times between steps [61] and times before unbinding [62]. This is also the most basic assumption we can make for times before binding.

#### A1.3.1 Stepping

Kinesin motors step processively along MT tracks in a hand-over-hand fashion, with each motor domain taking 16 nm steps [63] that move the center of mass of the motor forward by about 8 nm [64]. Unloaded, motors travel at velocity *v*_0_. When motors are subject to a hindering load, the velocity of the motor decreases with load until it nears 0 at the stall force, 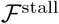. We use a form for this decrease studied in [65], where exponent *w* determines if the velocity decreases linearly, sub-linearly, or super-linearly. Asymmetrically, kinesin velocity is unaffected by assisting loads [66]. The overall form of the dependance is given by

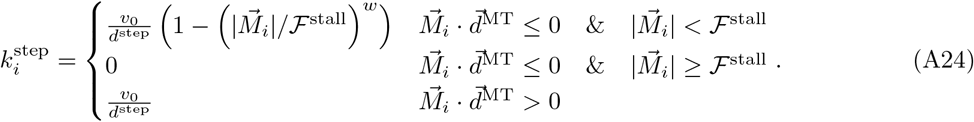

When a motor steps, it is moved forward along the direction of the MT to which it is bound by the step distance *d*^step^. This translates to an update of the head position

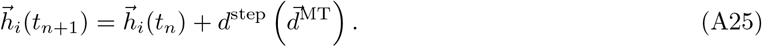

#### A1.3.2 Unbinding

Kinesin unbinds from the MT with a rate dependent on the force experienced by the motor. Hindering loads increase unbinding rate in an exponential fashion below the stall force [67]. Loads greater than the stall force continue to increase the unbinding rate, but do so more slowly, as found in [62] and analysis of data from [19] in [20]. For hindering loads, we use a piecewise exponential approximation of the theory of [20], in turn based on data from [19]. Unbinding rate is asymmetric, with assisting loads resulting in high unbinding rates [19, 68]. For assisting loads, we use an unbinding rate which is exponential with force, taken from the fit to data in [19]. Each of these exponential functions is parameterized as a function of force *F* by an unloaded rate *ϵ* and a characteristic force 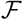, as 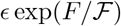. Together, we state the unbinding rate as

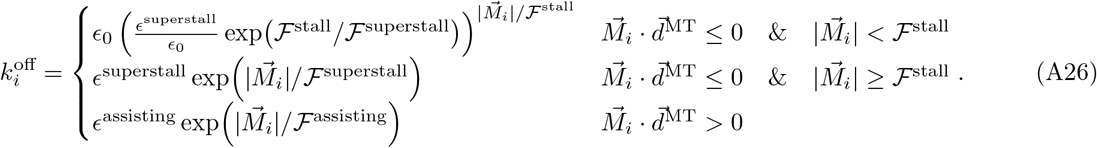

When a motor unbinds, it is simply put into the unbound state as defined at the beginning of this section.

#### A1.3.3 Binding

The conditions which govern the rate of binding to the MT are not well known. In the absence of detailed experimental elucidation, we make the assumption that the motor binds at a constant rate if the anchor is closer to the MT than its rest length. The probability of the motor binding at distances greater than the rest length decreases with distance proportionately to the probability of the motor spring stochastically taking on that distance. This translates to an on rate for motor *i* to the MT given by

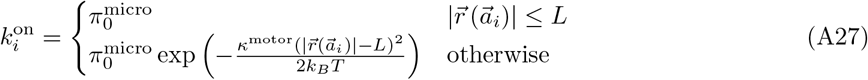

where 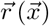 is given by equation A8. We note that we prevent the motor from binding when the motor stalk would intersect the cargo (if 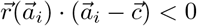), unless the motor is long enough to reach around the cargo to that point — that is, if

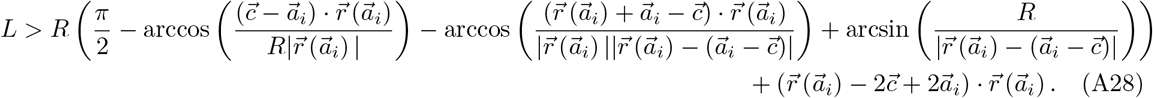

When a motor *i* binds to the MT, the head location is placed at the location on the MT nearest the anchor location 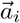, given by

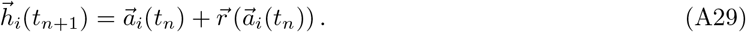

## A2 Numerical simulation of the model

Section A1 outlines a numerical scheme for updating for the model’s dynamic variables over a timestep Δ*t*. We simulate the model forward in time using these equations. Time steps are chosen dynamically. The largest stable time step for the Euler-Maruyama scheme is given by *ξ/κ*, where *ξ* is the drag coefficient and *κ* is the spring constant of the stiffest operating spring. The maximum time step is chosen based on the springs active during that step. The equivalent stiffness of multiple active motor springs is taken into account, but the steric spring in equation A7 remains by far stiffest in the system if it is active.

For each time step, we generate exponential random variables from distributions with means set by each Poisson rate, given in equations A24, A26, and A27, as in the Gillespie (next-event) algorithm. If any of the generated times are smaller than the maximum stable time step, the smallest generated time is chosen as the time step. If the chosen time came from an unbinding rate, a state change is implemented at the end of the update step by setting the motor to the unbound state. If this time came from a stepping rate or binding rate, the update occurs by equation A25 or A29, respectively. If no generated time is shorter than the maximum stable time step, the update is done with the maximum stable time step and there is no state change. Unlike in the Gillespie algorithm, rates are recalculated at each time step as they depend the changing locations and forces in the motors.

On average, many substeps are taken before a Poisson event occurs. The longest maximum stable time available to the system is the one characteristic of a single motor spring, which is ≈ 2.5 × 10^−5^ s. This is much shorter than the mean times between binding (≈ 0.1s), stepping (≈ 0.01s) and unbinding, even at several times the stall force (≈ 0.2 s at 20 pN). The shortest maximum stable time step normally experienced is that associated with the steric spring, which is ≈ 2 × 10^−7^ s, about 100 times shorter than the single motor maximum stable time step. During a run of the simulation, the cargo and MT overlap by an average of 2 nm. If the cargo and MT overlap by more than 5 nm for more than 5 consecutive time steps the simulation exits in an error.

The numerical simulation is written in C. It takes approximately 0.5 s to simulate 1s of cargo motion with a 3.3 GHz Intel Core i5 processor (single thread).

### A2.1 Model parameters

We attempt here to estimate the parameters relevant to cargos in the cell. We describe our estimations below. The parameters used to simulate the model are given in table A1.

#### A2.1.1 Motor parameters

We model our motors based on the well-studied kinesin-1 family. As such, many of the model parameters have been estimated in *in vitro* experiments. While rates of stepping and unbinding may be different in cells, the authors are not aware of any specific motor models from measurements in the cell, and it is not clear how individual motor behavior should be altered to reflect the cellular environment.

The length of the kinesin motor has been imaged in cryo-EM to be 80 nm [69, 70]. It has also been imaged attached to cargos [69, 71], where projections are shorter than 80 nm. It is unclear why. Some kinesin family members are shorter [72]. Since kinesin is floppy [73], molecules could also be folded. On the other hand, reach could also be extended by linker molecules. Since the length parameter represents the distance from the cargo kinesin can reach, not necessarily the most common length it assumes, we use 80 nm as the length parameter.

Motor binding rate has been measured in vitro to be about 5s^−1^ [74, 75]. It is known to be modulated by motor identity [75], presence of microtubule associated proteins such as Tau [47, 48] and MAP7 [41], and microtubule post-translational modification [76]. We have recently found that on rate is much higher for the first binding event than for subsequent events [5]. Since microtubule state is highly variable between transport scenarios and we do not know of any estimations of motor binding rate in the cell, we use the in vitro values of 100 s^−1^ for the first motor to bind the microtubule and 5 s^−1^ for subsequent motors.

#### A2.1.2 Cargo parameters

Cargos in the cell are highly variable in size, from neuronal vesicles on the order of 100nm [11, 18], to 3μm phagocytosed beads [2]. Other physical parameters, such as number of motors and motor diffusion coefficient are difficult to measure and not well understood. Neuronal vesicles have been measured to have just a few motors [11, 12], but it is unclear if other cargo types have similar numbers of motors, either in surface density or total number.

The surface viscosity of the cargo is discussed in Methods. It is largely unknown, therefore we explore a wide range of values in Results.

#### A2.1.3 Cytoplasm parameters

The interior of a eukaryotic cell is increasingly understood to be a complex environment with a complex rheology. Because it holds polymer networks such as those of actin and microtubules, along with numerous membrane-bound organelles, there are many obstacles to the movement of an object through the cytoplasm. The nature of this resistance is both viscous and elastic [21, 22]. Obstacles resist motion in a way that depends on the size and speed of the object being moved [22, 77]. In addition to being obstacles to motion, the cytoskeleton and organelles are active, generating non-thermal motion which pushes other objects in the cell [21, 22, 78]. Further large scale motion can be caused by movement of cell edges during cell motility [79].

In this paper, we seek to understand how molecular motors move cargos in this environment. The complexity of the effects described above, together with the fact that methods to mathematically describe the motion which results are under active development [80], necessitate that we simplify our description of the environment through which cargos are transported. In this paper we model cargos moving through, and diffusing in, a Newtonian fluid. While this description does not allow for elasticity, it does allow for more or less resistance to motion through changing the viscosity.

What viscosity best describes the cytoplasm? A wide range of values have been reported, spanning at least 5 orders of magnitude [49] from about ten times that of water [81] to over one million times [82]. Furthermore, there are a great many experiments — for example, Crick and Hughes estimated 1 Pa s in 1950 [83].

In [50], the authors use magnetic rotation microrheology on HeLa cells which have endocytosed magnetic nanoparticles. Endosomes in which these nanoparticles have gathered form a chain, which the authors rotate with magnets to probe the microrheology of the cytoplasm. The authors estimate a wide spread of viscosities for unperturbed HeLa cells, with a mean around 1 Pa s. However, when the HeLa cells are treated with nocodazole to disrupt microtubules or latrunculin A to perturb microtubules, they estimate a viscosity of 0. 4 Pa s, with a much tighter distribution of values. Since much of the motor attachment phenomena we describe in the paper is particularly influenced by cargo rotation, this study of rotating organelles is particularly relevant.

A similar study by Chevry et al. finds viscosities of 0.16Pas for NIH/3T3 cells and 1.4Pas for MDCK cells, using passive rotation of micron size wires [84]. Berret, using active rotations of similar wires in NIH/3T3 cells, finds viscosities around 30Pas for 2-3μm wires [85].

These three studies are able to learn about the rotation of an object in the cell by exploiting its elongated shape. One might think a spherical object might rotate more freely than an elongated one, as the elongated one could be pushing on other cellular components as it moves. Möller et al. estimate viscosity of J774 mouse macrophages using active rotations of micron size spherical magnetic particles and find viscosities between 10 Pa s and 60 Pa s [86]. As this is the only intracellular viscosity measurement made using spherical particles that the authors know of, we find no evidence to support a lower effective viscosity for spherical particles than for elongated ones. This fact also points out an opportunity for measurements of intracellular rheology using optical manipulation of spherical birefringent particles [87, 88] or tracking of janus particles [89].

Overall, we choose 0.4 Pa s as a representative value for cytoplasmic viscosity. Of the values measured for rotation in the papers listed above, it is toward the low end, being the same order of magnitude as the lowest value measured for rotation.

The cytoplasm also has many microtubules, and cargos can interact with multiple microtubules at once [16]. We limit the scope of this investigation to a single microtubule.

### A2.2 Initial conditions

**Anchor Placement** For cargos in the dispersed or free modes, we place anchors on the cargo by selecting points from a uniform random distribution over the surface of the sphere. For cargos in the clustered mode, all anchors are placed at the same point on the sphere.
**Cargo Orientation and Placement** Cargos are placed at a set distance from the MT in the +*z* direction and the cargo is aligned with the MT in the other two directions, i.e. (0,0, *R* + cargo-MT-dist + *R*^MT^). The orientation is such that the cargo north pole points in the +*z* direction. For simulations where motors pull cargos, one motor is chosen at random and the cargo is rotated so that motor is at the furthest point of the cargo in the — *z* direction. That motor begins bound to the MT. Cargo-MT distance is set to 0. For simulations where we measure time for the cargo to bind, the cargo is placed at a set cargo-MT distance and the orientation is not changed. No motors are bound.

## A3 Validation of the simulation

Simulations of 3D cargo motion coupled with the action of molecular motors have been previously reported [4, 16, 17, 60]. In this work we simulate cargos in which motor anchors experience drag through a viscous membrane and diffuse with a finite diffusion constant, which is novel. In this section, we show that our simulation agrees with analytical results in simplified situations.

### A3.1 Drag of anchors though the cargo membrane

In our model, anchors experience viscous drag when moving through the cargo membrane. While difficult to find for a spherical cargo geometry, an analytical solution for the anchor equation of motion is readily found for the case of a planar membrane. We begin with the sum of the forces on the anchor

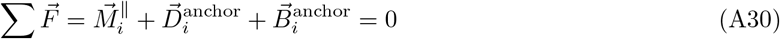

To test only the viscous drag, we set the brownian force 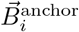 to 0. Inserting the forms of the motor force and viscous drag from equations A4 and A11, respectively, we simplify them for the 1D case of a static membrane. We note the motor head moves unidirectionally and restrict the solution to cases where *h* is constant in time and *h − a*(*t*) ≥ *L* (i.e. we start with the motor stretched) to obviate the need for a piecewise defined motor force. With these simplifications, equation A30 becomes

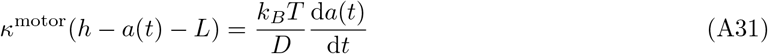

**Table A1:** List of parameters. Values listed are used for all simulations unless otherwise stated. *We note that with the form of equation A24 and these parameters, the force velocity curve is well matched to force-velocity data in [19].

| Parameter | Description | Value | Notes |
| --- | --- | --- | --- |
| $\kappa^{\text{motor}}$ | Motor spring constant | $320 \text{ pN } \mu\text{m}^{-1}$ | Measured in [57–59] |
| $L$ | Motor reach length | 80 nm | Length in electron microscopy, measured in [69, 70] |
| $v_0$ | Maximum velocity | $0.75 \text{ } \mu\text{m s}^{-1}$ | Measured in [19] |
| $d^{\text{step}}$ | Step size | 8.2 nm | Measured in [64, 90] |
| $w$ | Force-velocity curvature index | 1.8 | Measured in [91]* |
| $\mathcal{F}^{\text{stall}}$ | Stall force | 6.1 pN | Measured in [91]* |
| $\mathcal{F}^{\text{superstall}}$ | Characteristic detachment force for superstall loads | 7.6 pN | Fit to theory in [20], itself fit to data from [19] |
| $\mathcal{F}^{\text{assisting}}$ | Characteristic detachment force for assisting loads | 13 pN | Measured in [19]. |
| $\epsilon_0$ | 0-force unbinding rate | $0.8 \text{ s}^{-1}$ | Unloaded velocity divided by unloaded run length. Both quantities measured in [19] |
| $\epsilon^{\text{superstall}}$ | Base unbinding rate for superstall loads | $1.6 \text{ s}^{-1}$ | Fit to theory in [20], itself fit to data from [19] |
| $\epsilon^{\text{assisting}}$ | Base unbinding rate under assisting load | $7.4 \text{ s}^{-1}$ | Measured in [19] |
| $\pi_0^{\text{micro}}$ | Base binding rate (microscopic) | $100 \text{ s}^{-1}$ (first),<br>$5 \text{ s}^{-1}$ (subsequent) | Measured in [5],[74, 75]. |
| $N$ | Number of motors on the cargo | | Varied in simulations |
| $D$ | Diffusion constant of motor anchors in the cargo membrane | $0.1 \text{ } \mu\text{m}^2 \text{ s}^{-1}$ | Estimate for generic membrane bound protein from BioNumbers, BNID 114189 [10] |
| $R$ | Cargo radius | $0.25 \text{ } \mu\text{m}$ | Approximate size of lipid droplets in COS-1 cells [6] |
| $\eta$ | Viscosity of fluid immersing the cargo | $0.4 \text{ pN } \mu\text{m}^{-2} \text{ s}$ | Measured for active rotations of small vesicles in cells with a perturbed MT cytoskeleton in [50]. Measured for passive translational motion of tracer particles in [21]. |
| $k_B T$ | Thermal energy unit | $4.141 \text{ 95 pN nm}$ | at 300 K |
| $R^{\text{MT}}$ | MT radius | 12 nm | Measured in [92] |
| $\kappa^{\text{steric}}$ | Steric spring constant | $40 \text{ 000 pN } \mu\text{m}^{-1}$ | Set high enough to ensure cargo does not significantly intersect MT |
| $\vec{p}^{\text{MT}}$ | Defining point for MT | (0,0,0) | |
| $\vec{d}^{\text{MT}}$ | Defining direction vector for MT | (1,0,0) | |

**Figure A2:**
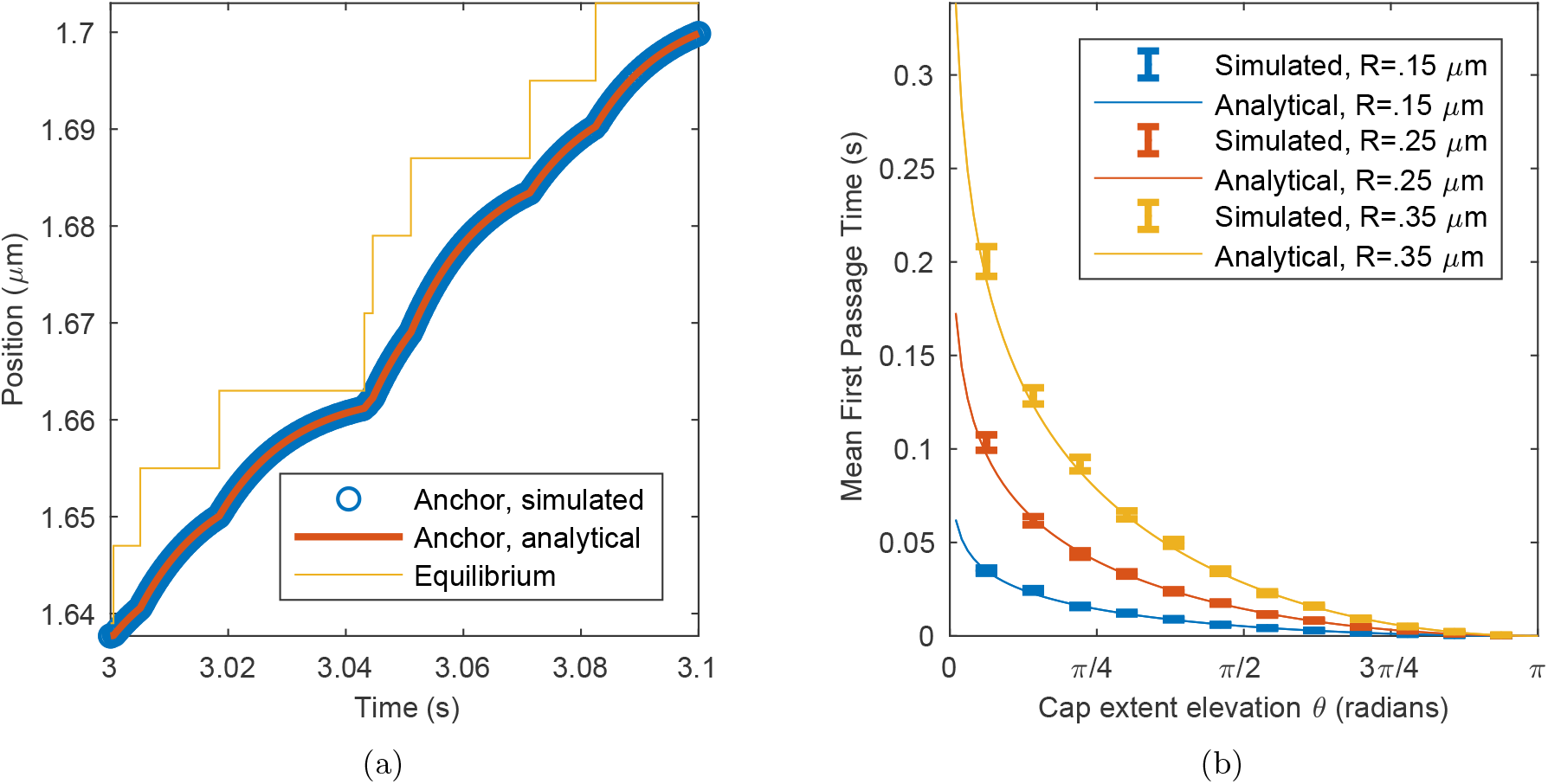
Comparison of simulated with analytical results for anchor drag and diffusion (a): Position of an anchor dragged through a 1D membrane by a stepping motor. A representative section of the time course of the anchor location in simulation (blue circles) is shown along with the equilibrium position of the motor spring (yellow). The analytical solution given this equilibrium location is found using equation A32 and shown in orange. Simulations done with *D* = 0.001 μm^2^ s^−1^. (b): Simulated and analytical mean first passage times for the anchor diffusing from the north pole of the cargo into a cap which extends up to elevation *θ*. For *θ* = 0, the cap is a point at the south pole. For *θ = π*/2, the cap is the southern hemisphere of the cargo. Theory is shown in solid curves, simulation data are shown as mean +/- SEM. Simulations done with *D* = 3 μm^2^ s^−1^.

Solving this differential equation with the initial condition *a*(0) = *a*_0_, we find

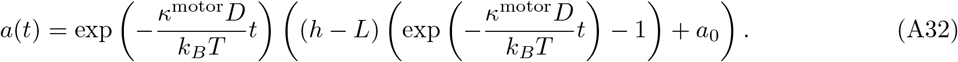

In the limit of a very large cargo with 0 cargo-MT distance, the simulation should produce this behavior. To compare the two, we first generate a simulation in this limit. The head location is constant between steps, allowing us to use the generated head locations to solve equation A32 in a piecewise manner over the entire time of the simulation. Analytical and simulated results are compared in figure A2a.

### A3.2 Anchor Diffusion in the Cargo Membrane

The mean first passage time for a particle to diffuse from the north pole of a sphere into a cap over the south pole which extends up to elevation *θ* is

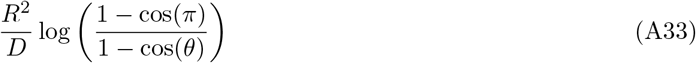

as given in [51]. To compare with this analytical solution, we simulate many trajectories of anchors diffusing from an initial location at the north pole of the cargo into caps extending to different elevations. The results of a comparison between the two is shown in figure A2b.

## Notes

### Competing Interest Statement

The authors have declared no competing interest.

### Summary of Updates

Update to title, and many edits to the main text for clarification. Also added link and DOI for github repository containing simulation code.

